# Identification of N-acetyl-rich heparan sulfate binding motifs and their role in expanding BMP distribution and signaling by Cerberus

**DOI:** 10.1101/2025.04.16.649141

**Authors:** Takayoshi Yamamoto, Yusuke Mii, Hidetoshi Saiga, Tatsuo Michiue, Masanori Taira

## Abstract

Secreted signaling molecules called morphogens including BMP and Wnt provide positional information by their concentration gradient. Although binding to an extracellular matrix, heparan sulfate (HS), is essential for morphogen gradient formation, it is still unclear how different ranges of gradients can be achieved. We previously found that two types of HS, N-acetyl-rich HS (NAc-HS) and N-sulfo-rich HS (NS-HS), are differentially clustered on the cell membrane, and bind to Frzb and Wnt8, respectively. Furthermore, we demonstrated that Frzb expands Wnt8 distribution by transferring Wnt8 from NS-HS to NAc-HS. This indicates that NAc-HS or its unsulfated form, heparan, which has been considered as a precursor of HS including N-sulfo HS, has an essential role in morphogen distribution. Here, we identified a so-called BMP antagonist, Cerberus (Cer), as another NAc-HS-binding protein. Binding of Cer to NAc-HS was enhanced by knockdown and repressed by overexpression of *ndst1*, which catalyzes the conversion of NAc-HS to NS-HS, suggesting that Cer binds to NAc-HS and/or heparan as does Frzb. We then explored NAc-HS binding motifs using deletion or alanine-substitution mutants of Frzb and Cer, and revealed that clusters of basic amino acids, Lys and Arg, are critical for their binding to NAc-HS. Functionally, Cer was able to expand distribution and signaling ranges of BMP4 through relocation of BMP4 from NS-HS to NAc-HS as has been shown with Wnt8 and Frzb. For this Cer activity, its NAc-HS binding motifs were essential. These results suggest that NAc-HS clusters and NAc-HS binding motifs constitute an interacting unit to modulate extracellular distribution of BMP4 as well as Wnt8.

## Introduction

It is conceivable that morphogens are secreted from source cells into surrounding extracellular space, and diffuse to form a concentration gradient, which orchestrates differentiation of many cell types to organize body patterning. Heparan sulfate (HS) and HS proteoglycans (HSPGs), such as glypicans and syndecans, have been shown to function as a platform to restrict distribution of secreted proteins (Yan and Lin, 2009), but how HSPGs contribute to distinct distribution ranges of morphogen remains to be unveiled.

HS belongs to the glycosaminoglycan (GAG) family and consists of repeating disaccharide units of N-acetylglucosamine (GlcNAc) and glucuronic acid (GlcA) with various modifications: epimerization of GlcA to Iduronic acid (IdoA), N-sulfonation (GlcNS), and 2-O-sulfation on GlcA/IdoA, and 6-O-and/or 3-O-sulfation on GlcNAc/GlcNS. HS has various combinations and degrees of modifications depending on tissue types as has been shown by disaccharide or oligosaccharide analyses of heparinase-digested or chemically cleaved HS extracts (Suflita et al., 2015). Based on these biochemical data (compositions of disaccharides with various modifications) and the assumption (HS chains expressed in each tissue are nearly homogenous), the following “NA and NS domain model” has long been postulated and seems to be widely accepted (Esko and Selleck, 2002; Sarrazin et al., 2011): that is, a single HS chain has a chimeric configuration with GlcNAc-rich regions (called NA domain), GlcNS-rich regions (NS domain), and alternating GlcNAc and GlcNS regions (NA/NS domain). However, it should be noted that any standard glycan sequencing methods have not been developed yet (Gray et al., 2019; Perez et al., 2023), and hence the “NA and NS domain model” has been still hypothetical We previously discovered that Wnt8 alone exhibits a short range of distribution and signaling as assayed by *Xenopus* gastrula embryos, but its short range of distribution and signaling could be expanded by secreted Wnt-binding proteins, Frzb (also known as sFRP3) and Crescent (Mii and Taira, 2009; Mii and Taira, 2011), both of which had been considered mere antagonists of Wnt. We have further demonstrated that HS chains with specific modifications have crucial roles in regulating Wnt distribution and signaling in *Xenopus* embryos (Mii, 2020; Mii et al., 2017).

HS on the core protein is initially synthesized as an unsulfated sugar chain of GlcNAc-GlcA disaccharide repeats, called heparan or a precursor of HS. Heparan in the cell may also be technically called N-acetyl-rich HS (referred to as NAc-HS) because heparan and NAc-HS are hard to be distinguishable in vivo with current experimental tools. Heparan is first modified by N-deacetyl/sulfotransferase (Ndst) to convert GlcNAc to N-sulfoglucosamine (GlcNS), generating N-sulfo-rich HS (NS-HS). NS-HS is further modified by other modification enzymes through stepwise processes, supposedly making various domains with different modifications in a single sugar chain (Esko and Selleck, 2002; Sarrazin et al., 2011; Yan and Lin, 2009). However, our cytological analysis of HS based on immunostaining revealed that at least two types of HS are discretely clustered on the cell membrane: one is NAc-HS clusters and the other is NS-HS clusters (Mii et al., 2017). We have also shown that the core protein of NAc-HS clusters is glypican 4 (Gpc4), an ortholog of *Drosophila* Dally-like, whereas the core protein of NS-HS clusters is either Gpc4 or glypican 5 (Gpc5), an ortholog of Dally (Mii et al., 2017). Furthermore, NS-HS clusters are frequently internalized compared to NAc-HS clusters, which tend to stay on the cell membrane. Importantly, based on these molecular features of HS clusters, we have discovered that these two types of HS clusters have different roles in modulating Wnt signaling. That is, Wnt8 preferentially associates with NS-HS clusters (Gpc4/5) and their complex is internalized to transduce Wnt signaling through signalosome formation, thereby leading to short distribution of Wnt8. By contrast, the secreted Wnt binding protein Frzb preferentially associates with NAc-HS clusters and hence stays in the extracellular space, leading to wide distribution of Frzb. Notably, the Frzb-Wnt8 complex associates with NAc-HS clusters (Gpc4), thereby relocating Wnt8 from NS-HS to NAc-HS clusters, inhibiting signalosome formation, and expanding Wnt8 distribution. These molecular mechanisms can explain how Frzb expands Wnt distribution and signaling ranges (Mii and Taira, 2009; Mii and Taira, 2011; Mii and Takada, 2020). Based on these results, we have proposed a new model that heparan (or NAc-HS) is not merely a precursor of NS-HS as well as other modified HS, but also functions as a binding platform for secreted “inhibitory” proteins, such as Frzb and most likely Crescent (Mii and Taira, 2009; Mii et al., 2017). This regulatory system of Wnts by their secreted binding proteins might be applicable to other morphogens, such as bone morphogenetic proteins (BMPs), but this possibility has not been tested yet.

BMP ligands belong to the transforming growth factor β (TGFβ) family, and play essential roles in dorsoventral patterning, proliferation and apoptosis in a concentration-dependent manner (Dosch et al., 1997; Hogan, 1996; Zakin and de Robertis, 2010). In *Xenopus* embryos, secreted antagonists for BMP have been identified as dorsalizing factors expressed in the Spemann-Mangold organizer region, such as Cerberus (Cer), Noggin (Nog), Follistatin (Fst), and Chordin (Chrd), which induce dorsal structures when ectopically expressed in the ventral region (De Robertis, 2009; De Robertis and Kuroda, 2004). In addition, Gremlin (Grem), which is expressed in neural crest cells in early *Xenopus* embryos, is also known to antagonize BMP (Hsu et al., 1998). Among antagonists of BMP, Chrd is known to shuttle BMP ligands (Bier and de Robertis, 2015; Inomata et al., 2013; Kuroda et al., 2004; Sasai et al., 1994; Zakin and de Robertis, 2010).

Cer is expressed in the anterior endoderm from the gastrula stage (Bouwmeester et al., 1996), and has an essential role in head formation (Kuroda et al., 2004). Although these antagonists have essential roles during embryogenesis, it is largely unknown how they are distributed from their source cells, except for Chrd and Sog (Ashe and Levine, 1999; Ben-Zvi et al., 2008; Srinivasan et al., 2002). In addition, the mechanism by which different distribution ranges are established among secreted molecules is still unclear.

To date, many “heparin-binding proteins” have been reported, such as antithrombin, peptide growth factors (FGF, Wnt, BMP, etc.), chemokines (IL8/CXCL8, SDF1α/CXCL12, etc.), some viral proteins (HSV gD, etc.) and so on (Ashikari-Hada et al., 2004; Esko and Selleck, 2002; Shriver et al., 2012). Heparin is a biological substance used biologically and medically as a blood anticoagulant because its highly sulfated pentasaccharide sequence binds and activates antithrombin (Izaguirre et al., 2014). Ligand binding sites in HS as well as HS binding regions/domains of ligands have been identified for antithrombin, FGF2, SDF1α/CXCL12, HSV gD, and so on (Ashikari-Hada et al., 2004; Esko and Selleck, 2002; Sarrazin et al., 2011), each of which specifically binds to a unique sequence of modified sugar residues containing 2OS, 6OS, and/or 3OS as well as IdoA. However, the molecular feature of NAc-HS binding domains remains largely unknown.

To address these unsolved subjects mentioned above, we first examined whether secreted BMP inhibitors bind to NAc-HS clusters and found that one of such binding proteins is Cer. Then, we analyzed and identified NAc-HS/heparan-binding motifs of Frzb and Cer. Finally, we attempted to generalize the Wnt8-Frzb regulatory system through NS-and NAc-HS clusters, focusing on the BMP4 and Cer regulatory system. We demonstrated that Cer expands distribution and signaling range of BMP4. Mutated Cer and Frzb lacking NAc-HS/heparan-binding motifs altered their abilities to modulate the distribution of BMP4 and Wnt8, respectively. Thus, this work highlights the importance of NAc-HS/heparan as a molecular basis for the differential distribution of BMP4 and its antagonists in the tissue patterning.

## Results

### Identification of N-acetyl-rich HS associating proteins

To identify NAc-HS associating proteins other than Frzb, we first examined secreted BMP antagonists, Cer, Grem, Nog, Fst and Chrd, focusing on whether their localization on the cell surface is affected by perturbation of *ndst1* expression using *Xenopus* embryos. We constructed plasmids for those proteins fused with monomeric Venus (mVenus/mV). mRNA was injected into the animal pole region of one ventral blastomere at the 4-cell stage, and *ndst1* mRNA or *ndst1* antisense morpholino oligos (MOs) into the other ventral blastomere as shown in a cartoon of Fig. 1a. We found that Cer-mV was detected in the intercellular space near the source cells, but significantly reduced on the cell surface of *ndst1*-expressing cells (Fig. 1a; *p*=1.5 x 10^-10^). Conversely, Cer-mV was significantly accumulated more on the cell surface of *ndst1*-knockdown cells than that of uninjected control (Fig. 1a; *p*=5.8 x 10^-11^). These data clearly showed that Cer-mV specifically associates with NAc-HS and/or heparan, not with NS-HS. Similarly, mV-tagged Grem, Nog, and weakly Fst prefers NAc-HS, but not NS-HS (Supplemental Fig. 1a, b). The preferential binding of Nog to NAc-HS appears to contradict its previously reported heparin-binding activity (Paine-Saunders et al., 2002), because heparin is rich in highly sulfated HS. However, it should be noted that heparin is a mixture of HS sugar chains with various degrees of sulfation, and hence “heparin-binding activity” does not necessarily mean “binding activity to highly sulfated HS.” This may be the case for the BMP antagonists according to our data, but it remains possible that they also bind to highly sulfated HS. Curiously, mV-Chrd was not detected in the intercellular space even near the source cells, implying that Chrd may not associate with extracellular matrices, similar to a secreted mVenus (Mii et al., 2021).

**Figure 1.**
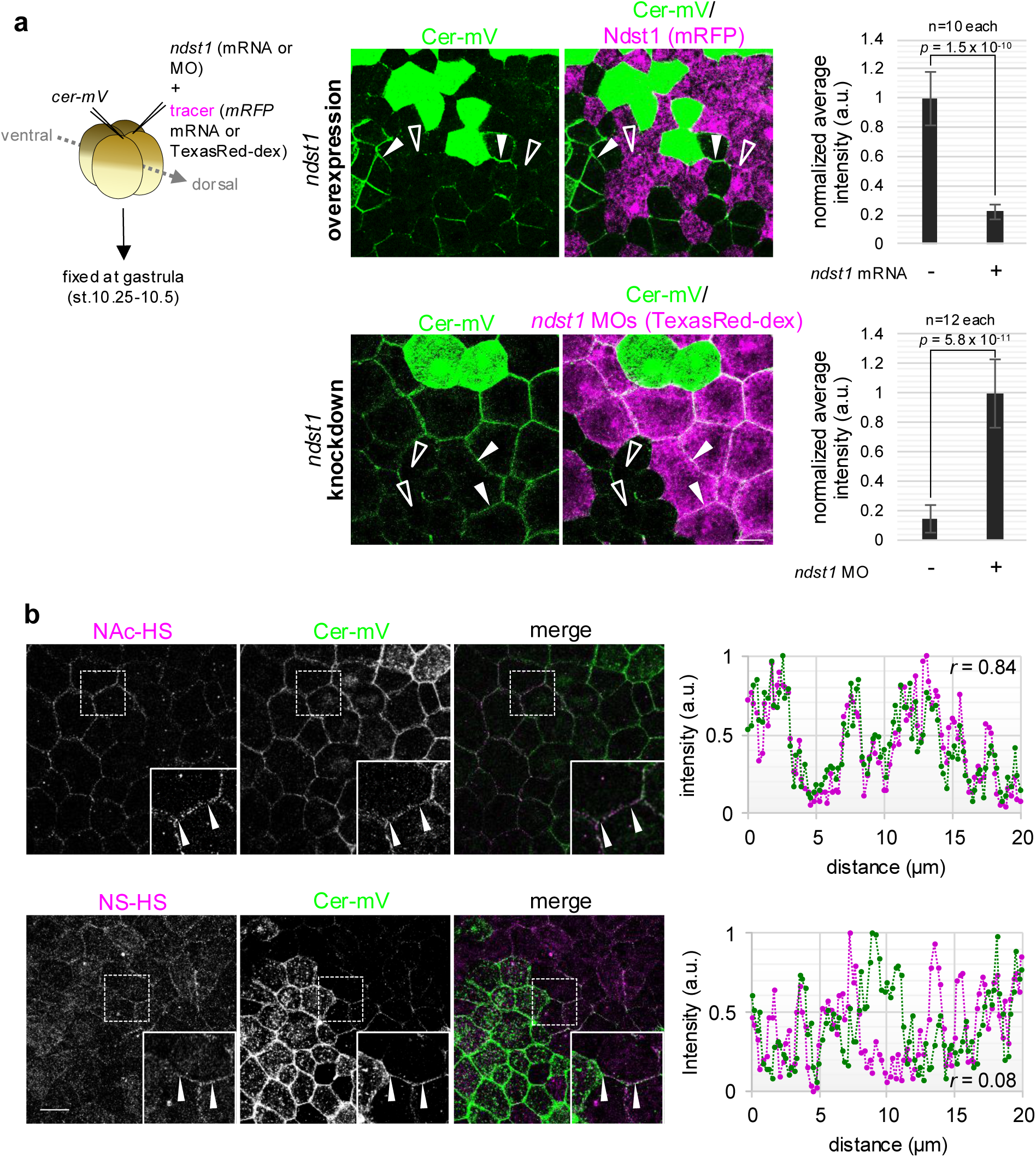
Cer localizes on N-acetyl-rich heparan sulfate (NAc-HS) a, Cer was well localized on the cells expressing lower level of *ndst1*. Cer mRNA and *ndst1* mRNA (with mRFP mRNA) or morpholino oligo (MO) (with Texas-Red dextran) were injected into different ventral blastomeres at the 4-cell stage. Cer-mV was not localized on the intercellular space between *ndst1*-overexpressing cells (open arrowhead), but on uninjected cells (arrowhead). Conversely, in the knockdown experiment, Cer-mV was localized on cells between the knocked-down cells (arrowhead). Graphs, quantification of Cer-mV intensity at cell boundaries of the knocked-down cells or uninjected cells (mean +/– s.e.m.; *p* = 1.5 x 10^-10^ (overexpression), *p* = 5.9 x 10^-11^ (knockdown), *t*-test). The number of measured cell boundaries is as indicated. b, Cer was localized on NAc-HS. Cer-mV was injected into the animal pole of a dorsal blastomere at the 4-cell stage and fixed at the gastrula stage (st.10.25-10.5). Specimens were processed to immunohistochemistry. Signal intensity was measured at region between arrowheads, and shown in graphs on the right. Values of correlation coefficients (*r*) are shown in the graphs. Amount of mRNA (pg/embryo): Cer-mV, 500 (a), 100 (b); Ndst1, 500; mRFP, 400. Amount of MOs (ng/embryo): *ndst1* MO, 14. Scale bars represent 30 μm (a-b).

To examine localization of the BMP antagonists to NAc-HS clusters by immunohistochemistry, we injected a small amount of mRNA for mV-tagged proteins, detected them with anti-GFP antibody, and visualized NAc-HS and NS-HS clusters with the specific antibodies, NAH46 and HepSS-1, respectively (Kure and Yoshie, 1986; Mii et al., 2017; Suzuki et al., 2008). As shown in Fig. 1b, Cer-mV were found to be well colocalized with NAc-HS clusters (r=0.84), whereas it was not colocalized with NS-HS clusters (r=0.08). Similarly, Grem-mV and mV-Nog were well colocalized with NAc-HS clusters (r=0.74 and 0.71, respectively; Supplemental Fig. 1c). These results suggest that Cer, Grem, and Nog, but not Chrd, are specifically associated with NAc-HS clusters on the cell membrane.

### Identification of NAc-HS-binding motifs of Cer and Frzb

To determine NAc-HS-binding regions and motifs, we prepared a series of Frzb deletion mutants fused mV. The constructs were expressed by mRNA injection, and their distributions in the intercellular space of *Xenopus* embryos were analyzed (Fig. 2a, Supplemental Fig. 2a). The data showed that when either the N-terminus or C-terminus of amino acid positions (aa) 273-311 was deleted, their intercellular distributions were dramatically reduced or barely detected (Fig. 2b, Supplemental Fig. 2b). As we have previously shown that a secreted HA-tagged GFP alone is not visible in the intercellular space outsides the source cells in *Xenopus* embryos by standard confocal microscopy (Mii et al., 2021), the loss of fluorescence signal in the intercellular space was considered as loss of binding ability of Frzb to NAc-HS. Therefore, we considered that the region of aa273-311, which contains 14 basic amino acids, is a NAc-HS binding region (Fig. 2a). We next examined distribution of Cer-mV and Cer-short (CerS)-mV, a N-terminally truncated form of Cer (aa110-270) (Piccolo et al., 1999) (Supplemental Fig. 3a). The data showed that CerS-mV was barely detectable in the intercellular space. Although a weak staining was still seen (Supplemental Fig. 3b), this was not reduced by Ndst1 (Supplemental Fig. 3c), indicating that the weak staining was not due to binding to NAc-HS. These observations suggest that the N-terminal sequence (aa1-109) of Cer contains an NAc-HS binding region.

**Figure 2.**
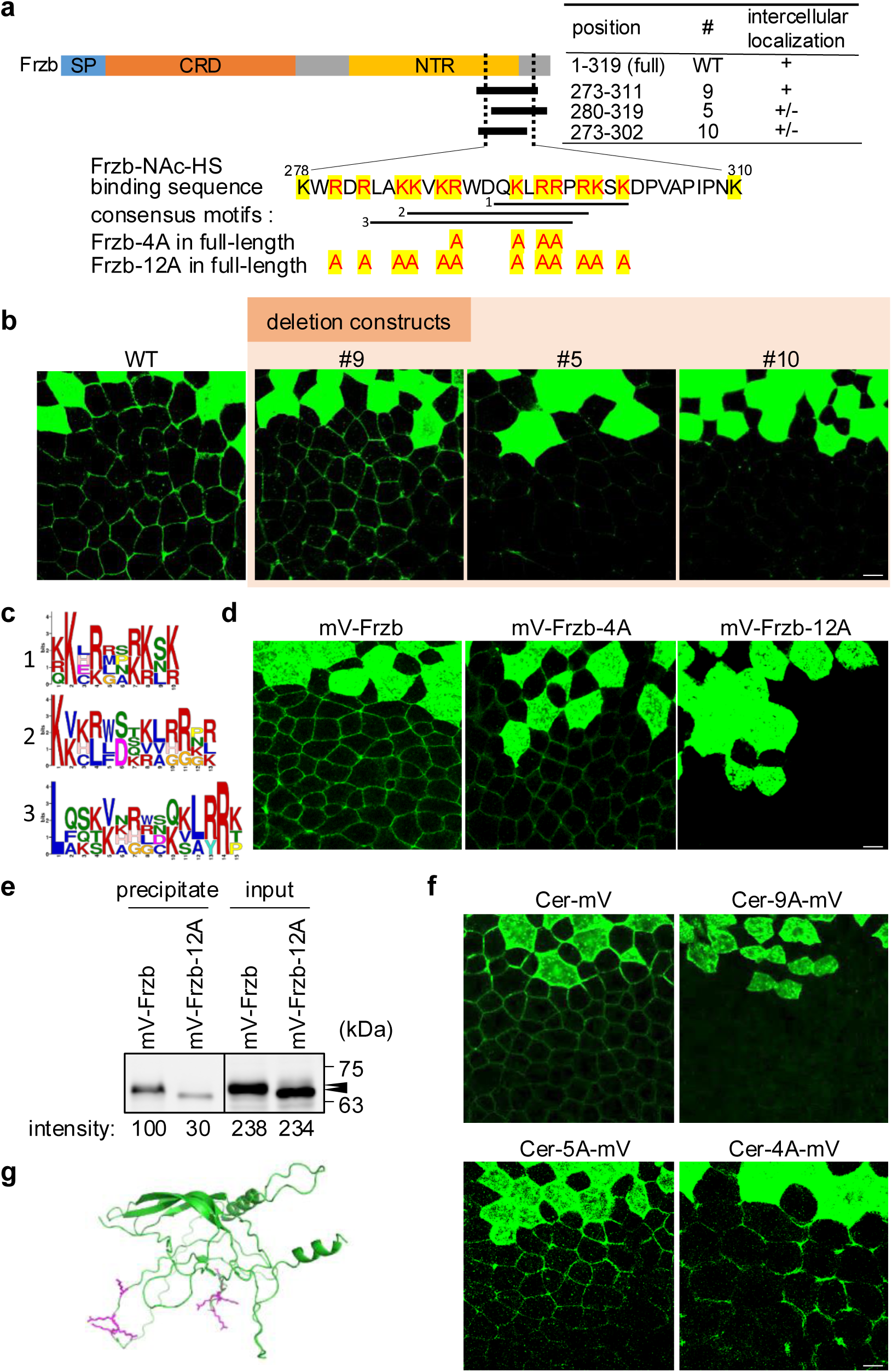
Identification of putative binding motifs to NAc-HS of Cer and Frzb a, Amino acid sequence of Frzb. Series of deletion constructs were indicated in black bars (positions of the sequences were indicated in amino acid number (start-end) at the right). Results of the intercellular localization (described in b) are summarized at the right. Sufficient sequence for association with the cellular membrane is between two dotted lines (black). The sequence is described below. Basic amino acids are highlighted in yellow. Residues changed to Ala are colored red. All series related to the experiment are described in Supplemental Fig. 2. b, Distribution of truncated constructs of Frzb. mRNAs of mV-tagged Frzb-del (deletion construct of Frzb) were injected into a blastomere at the 4-cell stage and observed at the early gastrula stage. c, Consensus sequence of Frzb (Lys-278 to Lys-310), CerN, Nog, Fst extracted by GLAM2 algorism (see Supplemental Information). d, Extracellular signals of mV-Frzb-12A were greatly reduced, compared with that of WT (mV-Frzb) and mV-Frzb-4A. mRNA of Frzb constructs was injected into a dorsal blastomere at the 4-cell stage and specimens were fixed at the early gastrula stage. e, Frzb bound to heparin by the putative NAc-HS-binding motif. mRNAs of mV-tagged Frzb constructs, were injected at the 4-cell stage. Cell extracts from stage 10.5 embryos were incubated with heparin beads, and the beads were washed with lysis buffer containing 100 mM NaCl. Results of all the series with the negative control are described in Supplemental Fig. 4b. The amount of protein bound to heparin beads, normalized by the mV-Frzb (100 mM NaCl) band, is indicated below each lane. f, Extracellular signals of Cer-9A-mV were greatly reduced, compared with the other constructs or WT (Cer-mV). mRNA of Cer constructs was injected into a dorsal blastomere at the 4-cell stage, and specimens were fixed at the early gastrula stage. g, Protein structure prediction of Cer with AlphaFold2 using ColabFold (Mirdita et al., 2022). Figure was described with PyMOL. Basic residues mutated to alanine in Cer-9A are colored magenta and the others are colored green. Amount of mRNA (pg/embryo): (b, d, and f) all mV-tagged Frzb or Cer constructs, 500. (e) all mV-tagged mRNA, 200. Scale bars represent 20 μm (b, d, f).

Using the GLAM2 algorithm, we searched for a consensus sequence among the minimal NAc-HS-binding region of Frzb, the N-terminal region of Cer, Nog and Fst, and obtained three motifs 1, 2, and 3, which are rich in basic residues (Fig. 2c). As shown in Fig. 2a, Frzb has an overlapping sequence of these three motifs in the NAc-HS binding sequence (Frzb-NAc-HS binding sequence). Examination of two alanine mutants of Frzb, Frzb-4A and Frzb-12A, showed that the intercellular distribution of Frzb-4A was partially reduced, whereas that of Frzb-12A was largely abolished (Fig. 2d). In vitro binding activity of the Frzb-NAc-HS binding sequence to heparan sulfate was further examined using heparin beads, which contain HS with various degrees of sulfation including highly sulfated HS and NAc-HS (Suflita et al., 2015). As expected, Frzb bound to heparin beads and Frzb-12A exhibited reduced binding ability (Fig. 2e, Supplemental Fig. 4a). This data suggests that the 12 Lys and Arg residues are necessary for direct binding to a component of heparin, most likely NAc-HS.

Cer has two separate regions with putative NAc-HS binding motifs 1 and 2 (Supplemental Fig. 3a; Cer-NAc-HS binding motifs 1 and 2). To examine them for NAc-HS binding ability, we constructed three alanine mutants of Cer, in which either of the two Lys/Arg motifs and both of them were substituted to alanine to make Cer-5A, Cer-4A, and Cer-9A, and examined their extracellular distributions. Extracellular signals of Cer-5A-mV or Cer-4A-mV were slightly reduced compared to Cer-mV, whereas those of Cer-9A-mV were greatly reduced (Fig. 2f), suggesting that the two regions containing Cer-NAc-HS binding motifs 1 and 2 redundantly bind to NAc-HS. This data is also consistent with the reduced extracellular distribution of CerS (Supplementary Fig. 3b), which lacks both Cer-NAc-HS binding motifs 1 and 2 (Supplementary Fig. 3a). The NAc-HS binding motifs of Cer are separated in the primary structure, which differs from those of Frzb. In the case of another CAN family protein Grem2, it has patches of basic residues, which are apparently divided into three separate blocks in the primary structure but are gathered in the 3D structure (Tatsinkam et al., 2015). Similarly, in a CAN family protein Sclerostin, the heparin binding residues are present on one side of the protein (Veverka et al., 2009). We performed protein structure prediction of Cer with AlphaFold2 and found that the basic residues of the motifs are on a single face of the structure (Fig. 2g). In addition, from the modelling of 3D structure of Frzb protein using SWISS-MODEL, a homology prediction workspace of protein structure, the motifs make α-helix, and the basic residues seem to be on a single face of the predicted α-helix (Supplemental Fig. 2c), which was also confirmed with helical wheel analysis (Supplemental Fig. 2d), although these analyses were held with a partial sequence of Frzb due to limitation of the homology prediction. To further confirm this, we performed protein structure prediction with whole protein sequence of Frzb using AlphaFold2, and this showed a similar result (Supplemental Fig. 2e). In addition, the NAc-HS/heparan binding motif of Frzb showed similarity to the reported heparin binding motif of human Nog protein (Paine-Saunders et al., 2002) (Supplemental Fig. 2f) and the basic residues of the Nog motif are also shown to be aligned on a single side of the α-helix (Paine-Saunders et al., 2002). These observations suggest that, regardless of whether the motifs are contiguous or separated, the motifs should be on a single side of the protein to bind to NAc-HS/heparan or heparin. Considering the distribution of secreted mVenus (Mii et al., 2021), it is possible that loss of NAc-HS binding ability of a protein will lead to expansion of its distribution range. To test this possibility, we used “morphotrap,” which is a membrane-tethered anti-GFP nanobody (Harmansa et al., 2015), so that unbound secreted mV-fusions can be detected. mRNA for mV-tagged Frzb or Cer constructs were injected into the animal pole region of one dorsal blastomere and *morphotrap* mRNA into two of the other blastomeres at the 4-cell stage (Supplemental Fig. 3d). We measured change of the signal intensity of mVenus along with distance from the source cells (see materials and methods for details and the legend for Supplemental Fig. 3e). As cells expressing morphotrap were not always contiguous and their expression levels were not identical, graphs with the cumulative mRFP signal intensity as X-axis were also constructed. As expected, the protein distribution of Frzb-12A and Cer-9A (blue lines) were expanded, compared to that of Frzb and Cer (red lines), respectively (Supplemental Fig. 3d). This indicates that loss of NAc-HS binding ability of a protein expands its distribution range.

We then examined whether the loss of NAc-HS-binding property of Frzb affects its ability to expand Wnt8 distribution. As previously reported (Mii and Taira, 2009), Frzb expanded Wnt8 distribution, but this ability was lost when NAc-HS binding motif was mutated (Supplemental Fig. 4b). Therefore, association of Frzb with NAc-HS is essential not only for its own distribution but also for that of its binding partner, Wnt8.

### BMP4 is associated with endogenous NS-HS and forms steep gradient

To test whether binding of the secreted antagonists to NAc-HS is important not only for Wnt but also for BMP signaling, we first examined how BMP4 is distributed in the intercellular space in *Xenopus* embryos. To address this, we made a construct, mVenus-tagged BMP4 (mV-BMP4) (Supplemental Fig. 5a), for visualization of BMP4 distribution. This construct showed ventralizing activity for *Xenopus* embryos when expressed dorsally, similar to untagged BMP4, suggesting that mV-BMP4 retains biological activity (Supplemental Fig. 5b). With this construct, we showed that fluorescence of mV-BMP4 was detected in a punctate manner in the intercellular spaces near the source cells (Fig. 3a, mV-BMP4). This short-range and punctate distribution in the intercellular space is reminiscent of that of Wnt8, which associates with NS-HS (Mii et al., 2017). To examine co-localization of BMP4 with NS-HS or NAc-HS, we injected a small amount of mV-BMP4 mRNA and stained simultaneously with anti-GFP antibody and anti-NS-HS or anti-NAc-HS antibody. Fig. 3b shows that BMP4 was well co-localized with NS-HS (*r* = 0.80), but not with NAc-HS (*r* = 0.06). This preferential association of BMP4 with NS-HS was confirmed by gain-of-and loss-of-function analyses for *ndst1* (Fig. 3c,d).

**Figure 3.**
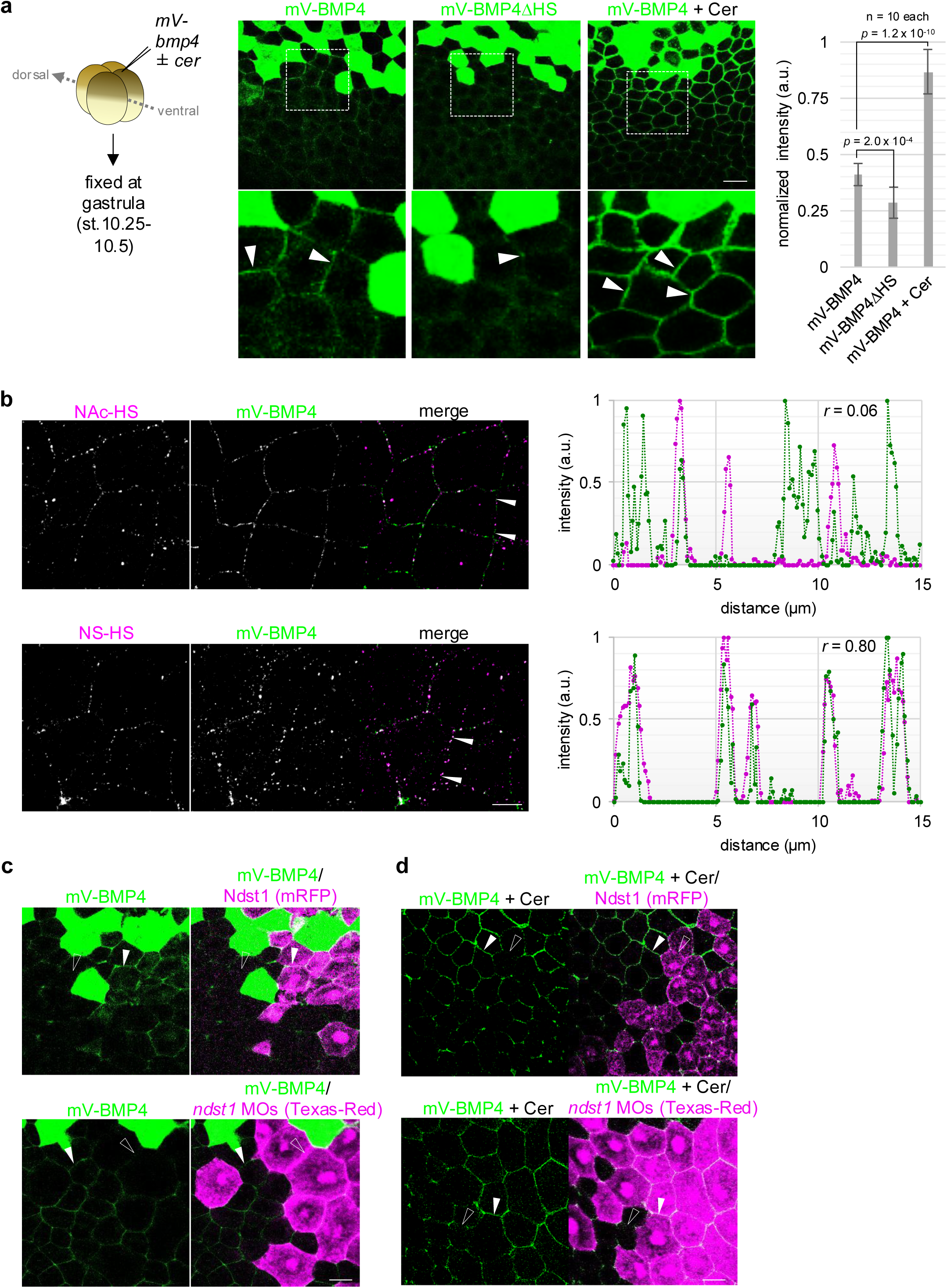
BMP4 is localized on N-sulfo-rich HS (NS-HS), and the distribution is expanded by Cerberus a, BMP4 distribution with or without putative HS-binding motif, containing many basic residues, in *Xenopus* embryos. mRNAs of mV-BMP4 constructs were injected into the animal pole region of a ventral blastomere at the 4-cell stage. Injected embryos were fixed at the early gastrula stage (st. 10.25-10.5). Green cells on the upper side are the source cells. Dotted areas in upper panels are magnified in lower panels. Signal intensity of mV-BMP4 in the intercellular region was higher than that of mV-BMP4ΔHS. The signal of BMP4 was strengthened and expanded by co-injection of Cer. Graph, quantification of mVenus intensity in the intercellular region (within two-cell distance from the source cells), normalized by the intensity at each source cell (mean +/– s.e.m., *t*-test). Number of measured cell-boundaries are as indicated. b, BMP4 was localized on NS-HS. mV-BMP4 was injected into the animal pole of a dorsal blastomere at the 4-cell stage, and fixed at the gastrula stage (st.10.25-10.5). Specimens were processed for immunohistochemistry. Signal intensity was measured at the region between arrowheads, and described in the graphs at the right. Values of the correlation coefficients (*r*) are shown on the upper side of the graphs. c, Localization of mV-BMP4 was increased by *ndst1* expression (arrowhead), and decreased by *ndst1* knockdown (openarrowhead). mV-BMP4 mRNA and *ndst1* (with mRFP) mRNA or its morpholino oligo (MO) (with Texas-Red dextran) were injected into different ventral blastomeres at the 4-cell stage. d, mV-BMP4 co-expressed with Cer was localized not on *ndst1*-expressing cells but on *ndst1*-knocked-down cells. mV-BMP4 mRNA with Cer mRNA and Ndst1 (with mRFP) mRNA or its MO (with Texas-Red dextran) were injected into different ventral blastomeres at the 4-cell stage. mV-BMP4 was localized on the intercellular space between *ndst1*-knocked-down cells (arrowhead), but not between on *ndst1*-expressing cells (open arrowhead). Amount of mRNA (pg/embryo): mVenus-tagged BMP4 constructs, 500; Cer constructs, 304 (equivalent mol. with mV-BMP4); dnBMPR, 500; mRFP, 400. Amount of MOs (ng/embryo): *ndst1* MO, 14. Scale bars represent 50 μm (a), 10 μm (b), 30μm (c, d).

Considering the previous report showing that BMP-activated area was expanded by loss of the HS-binding motif (BMP4ΔHS) (Ohkawara et al., 2002), it is expected that BMP4ΔHS widely spreads due to its loss of HS binding. To examine this, we made an mV-BMP4 mutant construct (mV-BMP4ΔHS), in which a putative HS binding motif was deleted, similar to BMP4ΔHS (Ohkawara et al., 2002) (Supplemental Fig. 5a). Fig. 3a shows that fluorescence intensity of mV-BMP4ΔHS was lower than that of mV-BMP4, even in the nearest location outsides the source cells. For quantitative comparison, we measured the fluorescence intensity of mV-BMP4 or mV-BMP4ΔHS in the intercellular regions surrounding non-source cells within a two-cell distance from the source cells. The results confirmed that the amount of mV-BMP4ΔHS was significantly lower than that of the control (mV-BMP4) (see Fig. 3a, right panel). To visualize a freely diffusing BMP4, we used a dominant-negative BMP receptor (dnBMPR) (Suzuki et al., 1994), in which the cytoplasmic domain is deleted. dnBMPR mRNA was injected into an animal blastomere away from a BMP4 mRNA-injected one at the 16-cell stage (Supplemental Fig. 6). mV-BMP4ΔHS, but not mV-BMP4, was detected on the cells expressing dnBMPR, even on the opposite side of the embryo (Supplemental Fig. 6). These results suggest that when BMP proteins do not bind to NS-HS, they increase the fraction unbound to cell surface, resulting in a wider distribution. It is also suggested that when BMPs bind to NS-HS, they are internalized with NS-HS, narrowing the distribution of BMPs.

### BMP4 distribution is expanded by Cer

We next examined whether the BMP antagonist Cer expands BMP4 distribution, as Frzb does for Wnt8. When Cer and BMP4 mRNAs were injected into the same blastomere or different blastomeres at the 4-cell stage, Cer dramatically expands BMP4 distribution over 7-cell distance (Fig. 3a, mV-BMP4 + Cer; Supplemental Fig. 7). The normalized signal intensity of BMP4 in the intercellular space near the source was increased, compared with that in the absence of Cer (Fig. 3a, right panel). Co-injection of various amounts of Cer with a fixed amount of BMP4 indicated that an equimolar amount of Cer was required for visualization of mV-BMP4 in the intercellular space near the source cells (Supplemental Fig. 8). When the splicing variant CerS, which lacks abilities binding to both NAc-HS (Supplemental Fig. 3a) and BMP4 (Piccolo et al., 1999), was co-expressed, the expansion of BMP4 distribution was not observed (Supplemental Fig. 7), suggesting that binding abilities of Cer to BMP4 and/or NAc-HS are essential for expanding BMP4 distribution. We then examined the requirement of the NAc-HS binding ability of Cer for the expansion of BMP4 distribution using Cer-9A-mV. As for BMP4 binding of Cer-9A-mV, Cer-mV and Cer-9A-mV exhibited a similar accumulation on membrane-anchored BMP (BMP4-GPI) expressing cells (Supplemental Fig. 3e), indicating that the absence of the two NAc-HS-binding motifs does not largely affect their BMP4 binding abilities. As shown in Fig. 4a, Cer-9A did not clearly accumulate mV-BMP4 on the intercellular space, compared with the wild-type Cer. These data suggest that the binding of Cer to NAc-HS is required for expanding distribution of BMP4 bound to the cell surface as the BMP-Cer complexes. The data also suggest that binding of the BMP-Cer complex to NAc-HS prevents BMP4 from internalization with NS-HS and the BMP receptor.

**Figure 4.**
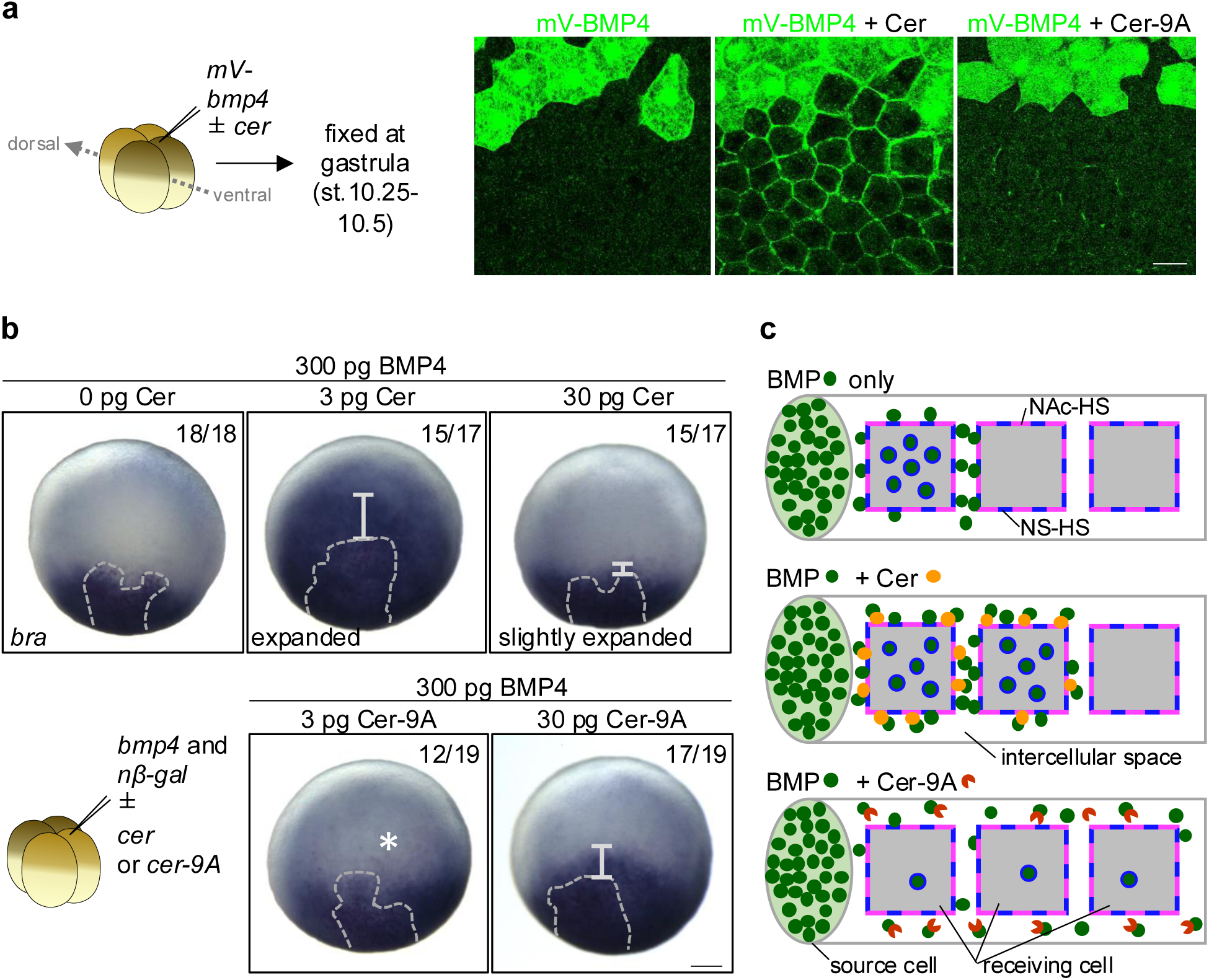
Expansion of BMP distribution and signaling range by Cer through binding to NAc-HS a, Expansion of mV-BMP4 distribution was observed with Cer, but not with Cer-9A under a standard confocal microscope. mV-BMP4 with/without Cer or Cer-9A was injected into a blastomere at the 4-cell stage. b, Area of *bra* expression induced by BMP4 injection was expanded by co-injection of Cer constructs. mRNAs of 300 pg BMP4 and n-*β*gal (tracer) with/without Cer or Cer-9A were injected into a dorsal blastomere at the 4-cell stage, and specimens were fixed at the gastrula stage (st.10.5). Source cells were colored red by Red-gal (surrounded by dashed lines). Activated area of BMP signaling is indicated by bars (chromogenic reactions were performed with the standard protocol; see materials and methods). c, Schematic figure of BMP expansion by Cer with NAc-HS. BMP binds to NS-HS on the cell membrane, and this complex internalizes into the cell with NS-HS as a sink. BMP in the cell means activation of BMP signal through binding to the receptor. Cer translocates BMP from NS-HS to NAc-HS as the BMP-Cer complex, and this prevents internalization of BMP, thereby expanding BMP distribution range. This also expands BMP signaling range (as indicated by number of BMP in the cells). Cer-9A prevents BMP from binding to the cell surface because Cer-9A no longer binds to NAc-HS on the cell membrane. Consequently, Cer-9A makes a shallow gradient of BMP as the BMP-Cer-9A complex. Note that this shallow gradient of BMP only very weakly activates BMP signaling. Thus, the steepness of the gradient of BMP signaling activity is BMP only > BMP + Cer > BMP + Cer-9A, as indicated the number of BMP in the cells. Amount of mRNA (pg/embryo): mV-BMP4, 500 (a), 300 (b); Cer constructs, 304 (equivalent mol. with mV-BMP4) in (a), 3 or 30 in (b). Scale bars represent 30 μm (a), 100 μm (b).

### Cer expands the signaling range of BMP

To see whether the expansion of “cell-surface-bound BMP” by Cer also expands the signaling range of BMP4, we examined the expression of a BMP target gene, *brachyury* (*bra*) by in situ hybridization when various amounts of Cer and a fixed amount of BMP4 were co-expressed by mRNA injection (Fig. 4b and Supplemental Fig. 9). When 3 pg Cer mRNA was injected, expansion of *bra* expression was observed. By contrast, 30 pg mRNA slightly expanded *bra* expression of similar strength (Fig. 4b) and 300 pg Cer mRNA injection almost eliminated *bra* expression from the entire region including the *bmp4/cer* source cells (Supplemental Fig. 9). Thus, Cer expanded *bra* expression (i.e., BMP4 signaling) in a concentration-dependent biphasic manner. This result is in contrast to the expansion of BMP4 distribution by Cer, i.e., the higher the amount of Cer, the higher the amount of BMP4 that accumulates near the source cells (Supplemental Fig. 8). These data suggest that a smaller amount of Cer expands both distribution and signaling of BMP4, whereas a higher amount of Cer exhibits more inhibitory effects on BMP4 signaling.

To test the requirement of the NAc-HS binding ability of Cer for expanding BMP signaling range, we carried out a similar experiment using Cer-9A instead of Cer (Fig. 4b). When 3 pg of Cer-9A mRNA was injected, very weak *bra* expression appeared to be observed almost throughout the embryo (Fig. 4b; white asterisk), which contrasts to the restricted expansion of much stronger *bra* expression by 3 pg Cer mRNA. When 30 pg mRNA was injected, Cer-9A resulted in expansion of relatively strong *bra* expression in comparison with Cer (Fig. 4b), which exhibited its inhibitory effects on BMP signaling. This implies that the inhibitory effect of Cer-9A on BMP4 signaling is weaker than that of Cer. Thus, it is suggested that NAc-HS binding of Cer is required to expand BMP4 signaling to its proper extent at its proper level in *Xenopus* embryos.

## Discussion

In this study, we aimed to reveal how morphogen distribution and signaling range are regulated. Our data suggest that Cer accumulates BMP4 in the intercellular space by binding to NAc-HS, thereby properly expanding the distribution and/or signaling range of BMP4 (Fig. 4c). Thus, Cer is capable of serving not only as an inhibitor against BMP signaling but also as an expander of BMP signaling range. In addition, this is the first report to identify NAc-HS/heparan binding motifs, thereby providing experimental evidence that NAc-HS binding of Cer and Frzb is important in the distribution and signaling of BMP4 and Wnt, respectively.

### Binding motifs of Cer and Frzb to NAc-HS

We have identified NAc-HS/heparan-binding motifs of Cer and Frzb. Although these motifs include similar sequences with five basic residues (noted as B) and 2 or 3 non-charged residues such as leucine, alanine, glycine and serine (noted as U) (BUUBBUBB or BBUBBUB or BBUBUUBB) (Supplemental Figs. 2a, 3a), the NAc-HS/heparan-binding motifs are hard to distinguish from heparin binding sequences. Many proteins that bind to heparin have been reported, and several binding motifs have been identified such as-XBBBXXBX-and –XBBXBX-, where B represents a basic residue and X a hydropathic residue (Cardin and Weintraub, 1989). However, any consensus sequence for heparin binding still remains to be elucidated (Rudd et al., 2017). This complexity in the binding motif is probably due to the fact that heparin and HS are both biological substances and hence they are mixtures of HS glycans with various degrees of modification (Chavaroche et al., 2013). In fact, Nog and Grem, which were identified in this study as NAc-binding proteins, have also been reported as heparin and/or HS-binding proteins (Paine-Saunders et al., 2002; Tatsinkam et al., 2015). Since we have revealed that the selective binding of morphogens and their secreted binding proteins to NAc-HS or NS-HS contributes the range of morphogen distribution, further analysis is required to understand the molecular basis of the selective binding to such HS glycans with specific modifications. For instance, identification of the difference in the binding motifs of NAc-HS and NS-HS is needed.

### NAc-HS and NS-HS are general modulators of the range of morphogen distribution

Several models for the distribution of secreted factors have been proposed. In “restricted diffusion model”, factors secreted from a source may bind to some extracellular matrices, such as heparan sulfate, which will interfere with their diffusion, and result in a steep gradient of secreted factors (Yan and Lin, 2009). In “source-sink model”, secreted factors, e.g. FGF, flow over the cell surface by free diffusion, meanwhile some of them may bind to a sink, such as their receptors, which will translocate the bound ligands into the cells. However, general mechanism for regulating different ranges of morphogen distribution remains to be elucidated. In the present study, BMP and its antagonist Cer bind to different modifications of HS; i.e., BMP binds to NS-HS whereas Cer binds to NAc-HS; and Cer with NAc-HS expands BMP distribution and signaling range, similar to the interaction between Wnt8 and Frzb via HS (Mii et al., 2017). Notably, many NAc-HS/heparan binding proteins have been discovered in this study, leading to the necessity of changing the perception that heparan is regarded as merely a precursor of HS. Taken together, we propose that not only sulfated HS (NS-HS), but also NAc-HS/heparan, are general modulators of morphogen distributions.

### Biological significance of Cer-mediated expansion of BMP signaling

Chrd distribution is reported to be detectable throughout almost the entire embryo by unconventional immunohistochemistry (Plouhinec et al., 2013), possibly reflecting its weak or no binding ability to the cell surface (Supplemental Fig. 1a). In addition, the necessity of long-range distribution of Chrd for the BMP shuttling is also suggested by mathematical modelling (Zinski et al., 2017). By contrast, Cer and/or the Cer-BMP4 complex bind to NAc-HS, thereby resulting in a more restricted spatial distribution of BMP4 compared to Cer-9A and Chrd. Moreover, because the core proteins, Gpc4 and Gpc5, of NAc-HS and NS-HS are localized on the lipid raft (Mii et al., 2017; Muñiz and Zurzolo, 2014), Cer on NAc-HS physically separates BMP from its receptors outside the lipid raft (Hartung et al., 2006) and from NS-HS clusters in the lipid raft. Thus, it is plausible that Cer restricts BMP signaling close to the *cer*-expressing cells (e.g. the anterior endomesoderm including the head organizer) by sequestering BMP4 to NAc-HS, which potentially leads to maintaining BMP pool and facilitating expansion of BMP4 distribution. These distinct molecular bases of Chrd and Cer suggest a complementary regulation of BMP gradients, supporting the reported synergistic effect on head formation (Kuroda et al., 2004). Furthermore, it is speculated that Cer expands not only BMP but also Wnt and Nodal signaling in a proper range around *cer*-expressing cells, which may also contribute to head formation. Local accumulation of ligands such as BMP4 on NAc-HS by Cer, along with signal inhibition, is thought to be crucial for precise patterning. Thus, this study leads to better understanding of regulation of morphogens during development and may provide a new insight into regeneration and clinical issues mediated by morphogens.

## Material and Methods

### *Xenopus* embryo manipulation and microinjection

All animal experiments were approved by the Office for Life Science Research Ethics and Safety, at the University of Tokyo. Manipulation of *X. laevis* embryos and microinjection experiments were carried out according to standard methods as previously described (Mii and Taira, 2009; Sive et al., 2000). Briefly, unfertilized eggs were obtained from female frogs injected with gonadotropin (ASKA Pharmaceutical), and artificially fertilized with testis homogenate. Fertilized eggs were dejellied with 2% L-cysteine-HCl solution (pH7.8) and incubated in 1/10x Steinberg’s solution at 14-20°C. Embryos were staged according to Nieuwkoop and Faber (Nieuwkoop and Faber, 1967). Amounts of injected mRNAs are described in the figure legends. For tracing MO-injected cells, 2.5 ng of Texas-Red dextran (Molecular Probes, D-1863) was co-injected.

### Morpholino oligos (MOs)

Because *Xenopus laevis* is an allotetraploid, we used a 1:1 mixture of two MOs targeting transcripts from both homeologs as follows: *ndst1*, AGGAATGGCACAAGCTCACAAATGC and AGGAGTGGCACAAGCTCACAAATGC (Mii et al., 2017).

### Plasmids and mRNA synthesis

pCSf107mT (Mii and Taira, 2009) was used as a vector possessing 4xSP6/T7 transcription terminator sequences to make plasmid constructs for mRNA synthesis unless otherwise mentioned. mRNAs were transcribed in vitro using an mMessage mMachine SP6 kit (Thermo).

For BMP4 constructs, tags (mVenus or Myc) were inserted into the BMP4 sequence after its cleavage site as previously reported (Degnin et al., 2004). Three fragments (N-terminus, tags and C-terminus) were independently amplified using the primers described in Supplemental Table 1. These products were inserted into pCSf107mT that was digested by BamHI and XbaI beforehand using In-Fusion system (Clontech).

pCSf107-Cer-mV-T was constructed by inserting a PCR product for the CDS using the primers described in Supplemental Table 1 into pCSf107mV-T using the BamHI/AvaI sites. pCSf107-Cer-5A-mV-T, Cer-4A-mV-T, Cer-9A-mV-T were constructed by inserting point mutations with PCR-based method into pCSf107-Cer-mV-T using the primers described in Supplemental Table 1 and universal primers (SP6 or T7).

N-terminus deletion constructs of mV-Frzb were constructed by inserting various regions of the Frzb sequence (amplified from pCSf107-Frzb-mV-T (Mii and Taira, 2009) using the primers described in Supplemental Table 1 as a forward primer and pCS2-R1 primer as a reverse primer) into pCSf107mVmT vector using the ClaI/XbaI sites. Similarly, C-terminus deletion constructs of mV-Frzb were amplified from pCSf107-Frzb-47AA-mV-T using eGFP-F1 as a forward primer and the primers described in Supplemental Table 1 as a reverse primer. pCSf107-Ndst1-T (Mii et al., 2017) was used as a template for expressing Ndst1. To make myc-BMP4-GPI construct, myc-BMP4 was amplified from pCSf107-myc-BMP4 using the primers, SP6CF1+ and BMP4GPI-R (Supplemental Table 1), and it was inserted into pCSf107mGPI-T vector (Mii et al., 2017) using the BamHI/XbaI sites. For mV-Chrd, mV-Nog and mV-Fst constructs, the CDSs were amplified from cDNA of *X. laevis* (J. strain) by the primers described in Supplemental table 1, and these were inserted into pCSf107mV-mT by EcoRI/XbaI (BamHI/XbaI for Fst) sites.

### Immunohistochemistry and antibodies

Embryos were fixed with MEMFA (0.1 M MOPS, pH 7.4, 2 mM EGTA, 1 mM MgSO4, 3.7% formaldehyde) and immunostained by standard protocols with Tris-buffered saline (Shibata et al., 2005; Sive et al., 2000). For permeabilization, 0.1% Triton X-100 was used. Primary antibodies used for immunostaining were as follows: mouse monoclonal anti-Myc (9E10, BIOMOL, 1:1000), anti-Myc (4A6, Upstate, 1:2000), anti-NAc-HS (NAH46, Seikagaku, 1:200), anti-NS-HS (HepSS-1, Seikagaku, 1:200); anti-pSmad1/5/9 (13820, Cell signaling technology, 1:2000). Secondary antibodies used were goat or donkey polyclonal anti-mouse or rabbit IgM or IgG antibodies labelled with Alexa Fluor 488, Alexa Fluor 546, Alexa Fluor 555 or Alexa Fluor 647 (1:500, Molecular Probes). The specimens were visualized by the confocal microscope (LSM710 (Zeiss) or FV-1200 (Olympus)). The signal intensity was measured by Fiji software (ImageJ 1.53f51; Java 1.8.0_172 (64-bit)) (Schindelin et al., 2012).

### Whole mount in situ hybridization

Whole mount in situ hybridization (WISH) was performed based on *Xenopus* standard method (Harland, 1991) with slight modifications in the duration of washes and hybridization temperature of 65°C. DIG labelling was performed using T7 RNA polymerases and DIG labelling mix (Roche, Mannheim, Germany).

### Heparin-binding assay

The binding of Frzb constructs to heparin was analyzed using Affi-Gel® Heparin Gel (Bio-Rad Laboratories). Nine embryos injected with mRNA for mV-tagged Frzb constructs were lysed in 90 μL of lysis buffer (10 mM Tris–HCl (pH 7.5), 100 mM NaCl, 5 mM EDTA, 0.5% Nonidet P-40, protease inhibitor cocktail (Complete EDTA-free, Roche)) with pipetting. The cell lysates were centrifuged at 15 kG. 20 μL of the supernatant was incubated with 20 μL of heparin beads containing 190 μL of the lysis buffer for 2 h at 4°C. The heparin beads were washed three times with lysis buffer containing 100 mM to 800 mM of NaCl, and boiled in SDS sample buffer at 40°C for 30 min and 98°C for 5 min. Immunoblotting was performed as described (Hara et al., 2012). mV-tagged proteins were analyzed with GFP Antibody IRDye® 800 Conjugated (600-132-215, Rockland Immunochemicals, Inc.).

## Acknowledgement

We thank Dr. Takafumi Ikeda for critical comments on this project, Dr. Yuta Otsuka for help in starting this work and critical comments on this project, Dr. Takehiko Nakamura (Seikagaku corp., Japan) for NAH46 antibody and hybridoma, Dr. Osamu Yoshie (Kindai University, Japan) for HepSS1 antibody, Dr. Makoto Matsuyama (Shigei Medical Research Institute, Japan) for the contribution to the generation of NAH46 and HepSS-1 antibody from the hybridomas, Dr. Edward M De Robertis (University of California, USA) for pCS2+Cer, Dr. Ira Blitz (University of California, USA) for Tsg construct, Dr. Hiroki Kuroda (Keio University, Japan) for Cer-short construct, Dr. Jim Smith (National Institute for Medical Research, UK) for Brachyury construct, Dr. Naoto Ueno (NIBB, Japan) for mouse BMP receptor dominant negative mutant (dnBMPR), and Dr. Hiroyuki Takeda (University of Tokyo, Japan) for Zeiss LSM710 confocal microscope. This work was supported in part by MEXT/JSPS KAKENHI (25251026, 18H02447, and 21K06126 to MT, 19K16138 to TY, 18K06244, 21K06183, and 23K05791 to TY and TM), Narishige Zoological Science Award (to TY) and Daiwa Anglo-Japanese foundation (to TY). We also thank National BioResource Project (NBRP) and NBRP Information Center (National Institute for Genetics) for providing the *Xenopus* genomic database (http://viewer.shigen.info/xenopus/).

## Author contributions

TY, YM, and MT conceived this project; TY performed experiments; TY, YM, HS, and MT wrote the original draft; TY, YM, HS, and MT reviewed and edited the manuscript.

## Competing interests

The authors declare no competing interests.

## Supplemental information

**Supplemental Figure 1.**
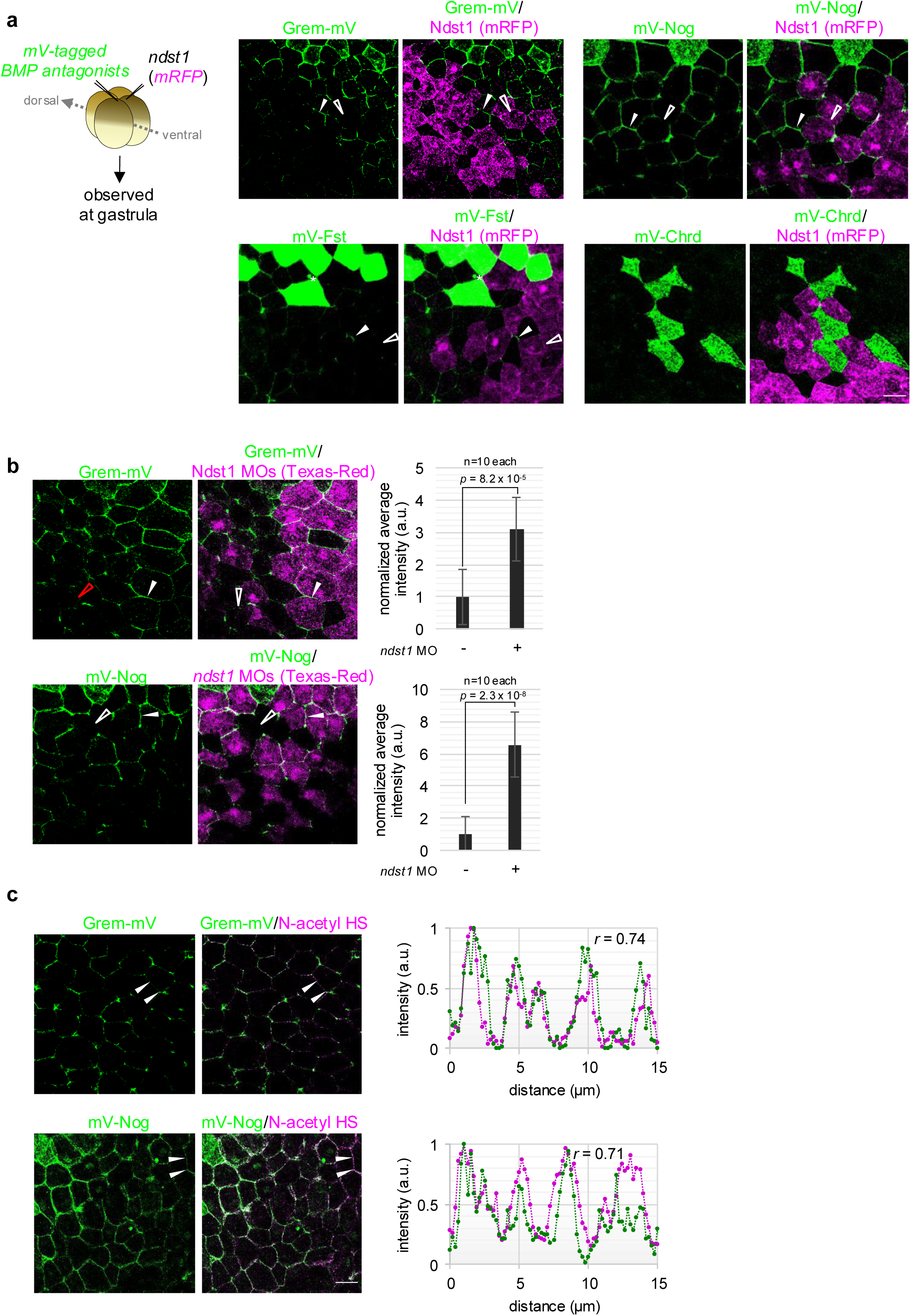
Distribution of various BMP antagonists and its change by Ndst1 expression. a, To examine difference in distribution of the antagonists, we injected mRNAs of mVenus-tagged antagonist and Ndst1 (with mRFP) were injected into dorsal different blastomeres at the 4-cell stage. Source cells were colored green even in the intercellular region. Although distribution of Chrd was not clearly observed in the intercellular region, those of Cer, Nog, Fst and Grem were in the intercellular region (arrowhead). In addition, Grem, Nog, and Fst were not localized on Ndst1-expressing cells (open arrowhead), suggesting these proteins are NAc-HS associating proteins. b, Grem and Nog were localized on the cells between *ndst1*-knocked-down cells (arrowhead). Graphs, quantification of mVenus intensity at cell boundaries of *ndst1*-knocked-down cells or uninjected cells (mean +/– s.e.m.; *t*-test). Number of measured cell boundaries are as indicated. c, Grem and Nog were localized on NAc-HS. mV-tagged Grem or mV-tagged Nog was injected into the animal pole of a dorsal blastomere at the 4-cell stage, and fixed at the gastrula stage (st.10.25-10.5). Specimens were processed for immunohistochemistry. Signal intensity was measured at the region between arrowheads, and described in the graphs at the right. Values of the correlation coefficients (*r*) are shown in the graphs. Amount of mRNA (pg/embryo): mVenus-tagged antagonists, 500 (a,b), 100 (c); Ndst1, 500; mRFP, 400. Amount of MO (ng/embryo): *ndst1* MO, 14. Scale bar represents 30 μm.

**Supplemental Figure 2.**
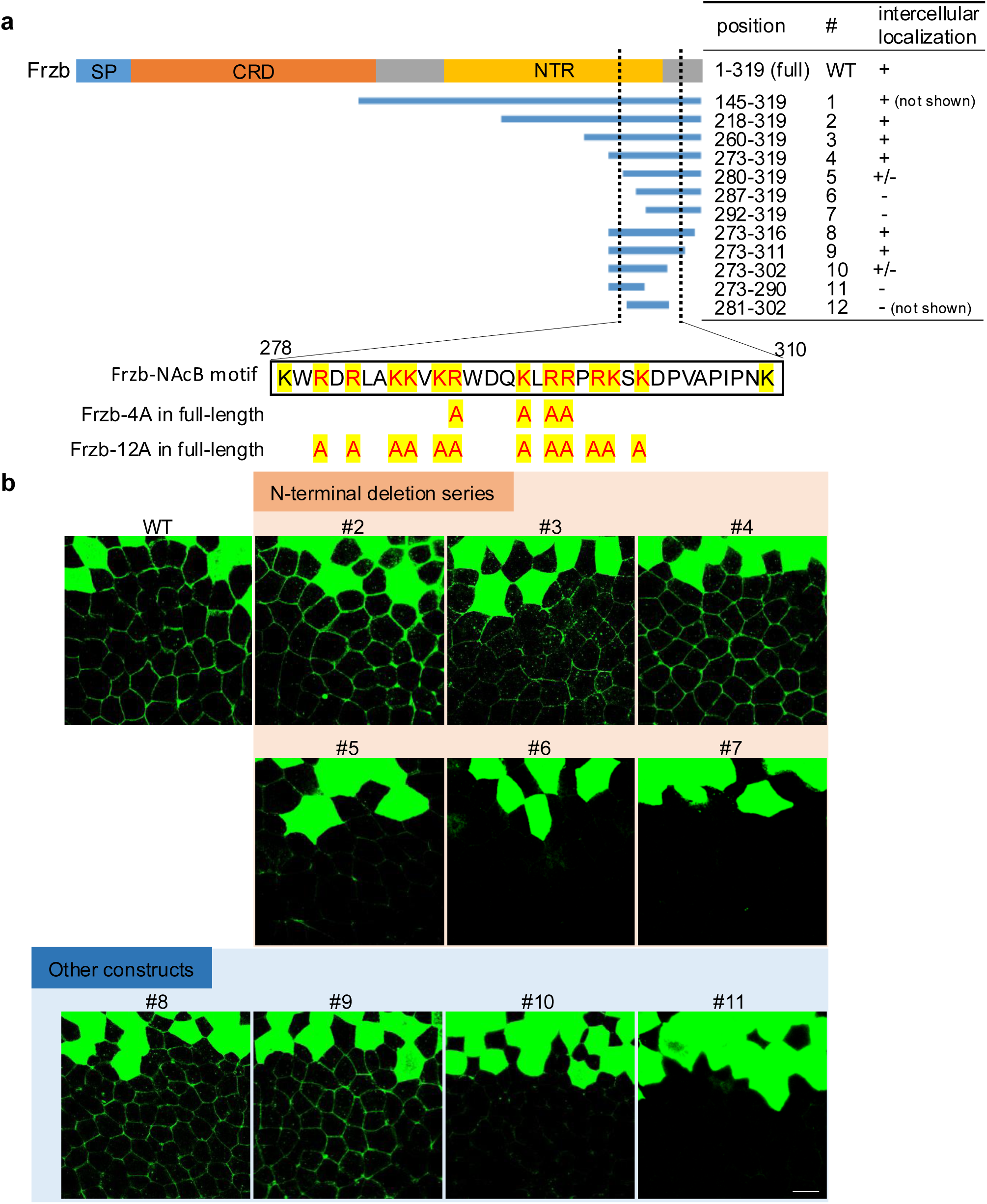
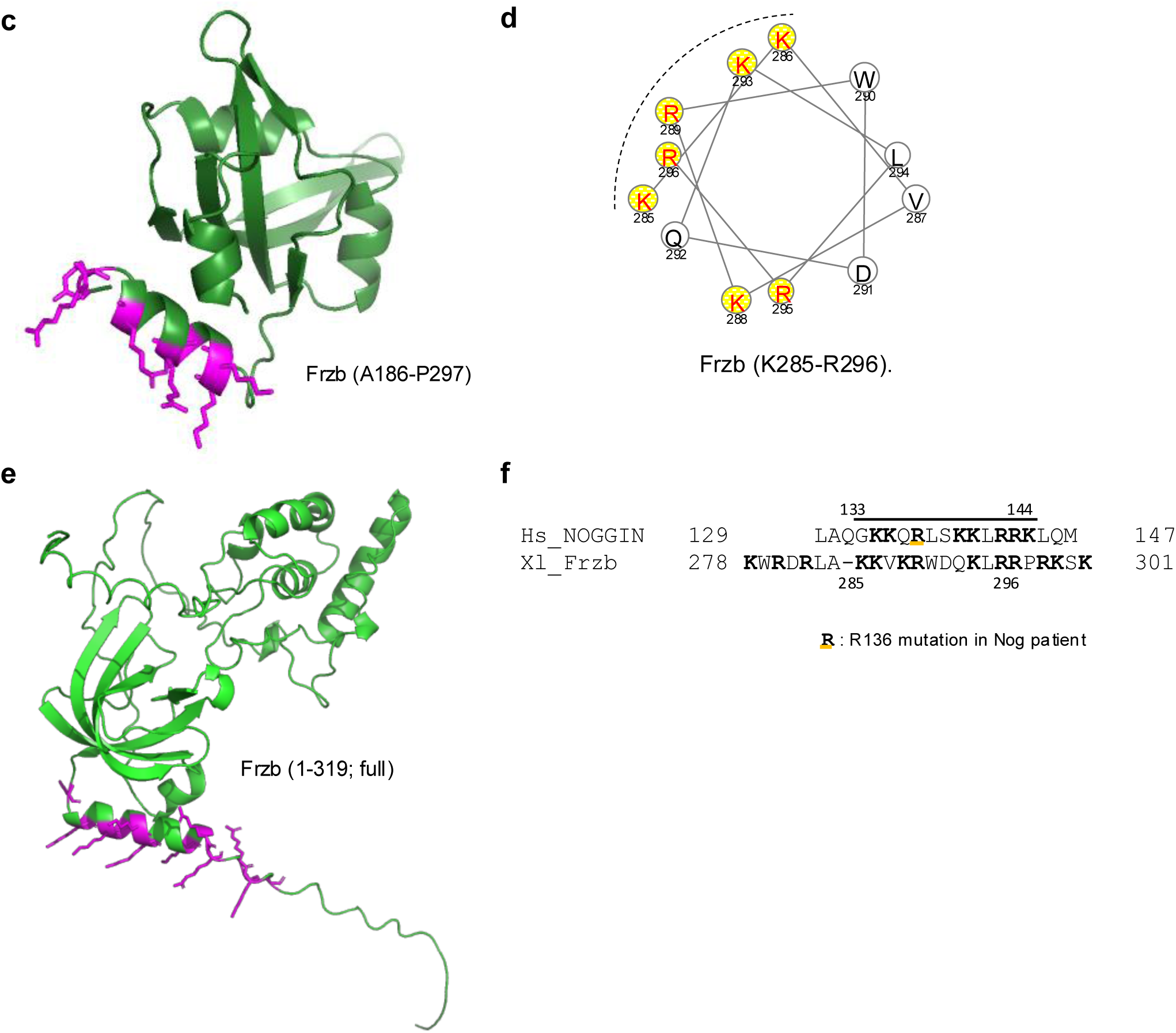
Considerations on the binding motif of Frzb to N-acetyl-rich HS/heparan. **a**, Amino acid sequence of Frzb. Series of deletion constructs were indicated in blue bars Positions of the sequences were indicated in amino acid number (start-end) at the right. Results of the intercellular localization (described in b) are summarized at the right. Sufficient sequence for association with the cellular membrane is between two dotted lines (black). The sequence of region 278-310 and the positions of Ala mutations are shown below. Basic amino acids are highlighted in yellow. Residues changed to Ala are colored in red. **b,** Distribution of truncated variants of Frzb. mRNAs of mV-tagged Frzb-del (deletion construct of Frzb) were injected into a blastomere at the 4-cell stage and observed at the early gastrula stage. Some of these data are also described in Figure 2. **c,** Protein structure prediction of partial sequence of Frzb (A186-P297) are built using SWISS-MODEL (https://swissmodel.expasy.org) based on the structure of a secreted Frizzled-related protein, sizzled (PDB code 5xgp). Figure was described with PyMOL. Basic residues mutated to alanine in Frzb-12A are colored magenta and the others are colored green. **d,** Helical wheel analysis of the sequence predicted to make α-helix in the putative NAc-HS binding motif of Frzb in **c** (K285-R296). Basic residues mutated to alanine in Frzb-12A seem to be on a single face of the predicted α-helix (dashed line). **e,** Protein structure prediction of Frzb with AlphaFold2 using ColabFold (Mirdita et al., 2022). Figure was described with PyMOL. Basic residues mutated to alanine in Cer-9A are colored magenta and the others are colored green. **f,** Partial alignment of human Noggin (Nog) and Xenopus Frzb in heparan sulfate binding region. Nog (aa129-147) and Frzb (aa278-301; a sufficient sequence for NAc-HS/heparan binding). Note that A Noggin deletion mutant, hNogΔB2 (Δ133-144; lined), does not bind to heparin (Paine-Saunders et al., 2002). The residue (underlined “R”) of Nog mutated in dominant disorders proximal symphalangism was underlined (Masuda et al., 2014). Amount of mRNA (pg/embryo): mVenus-tagged Frzb variants, 500. Scale bar represents 30 μm.

**Supplemental Figure 3.**
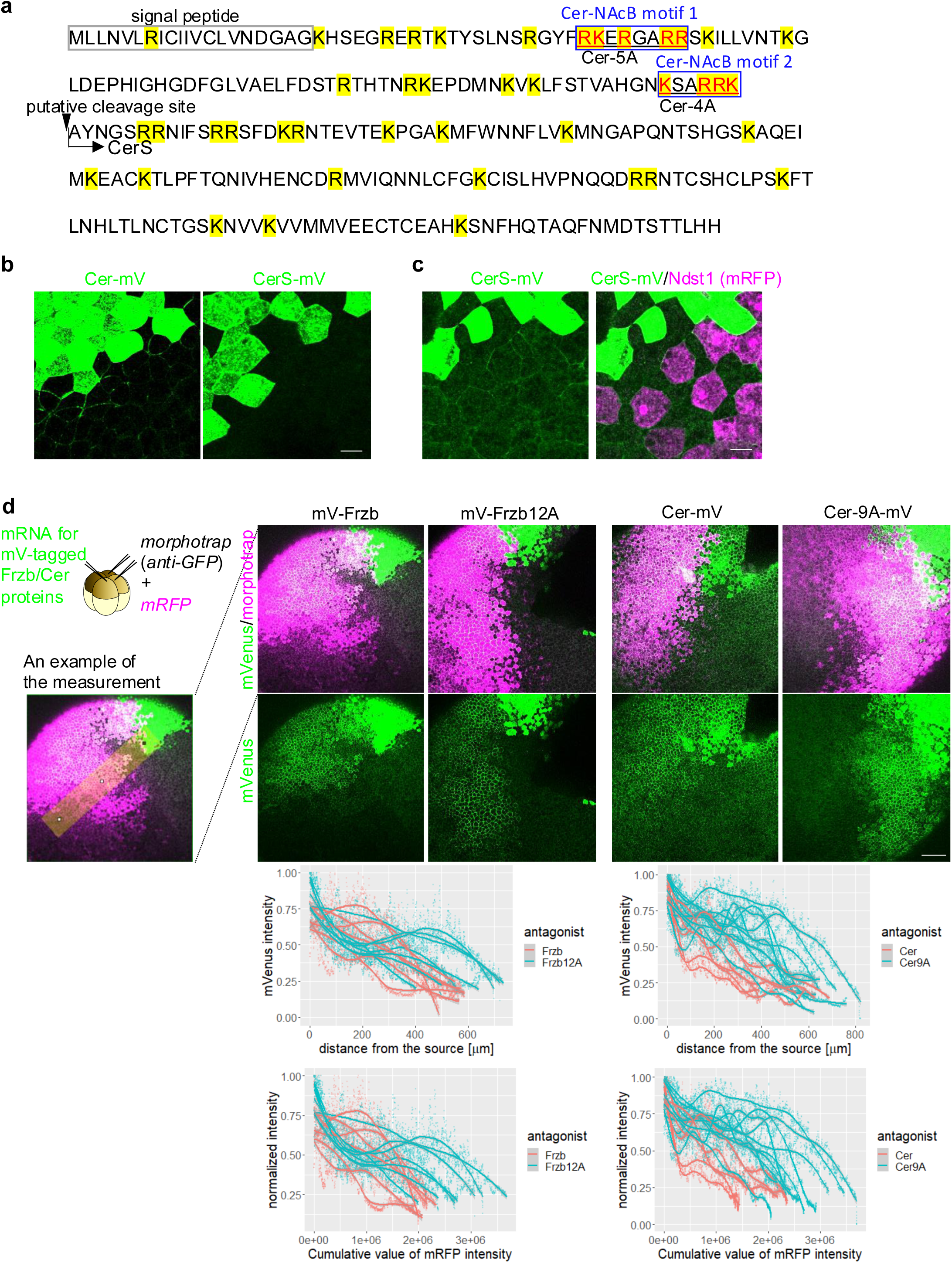
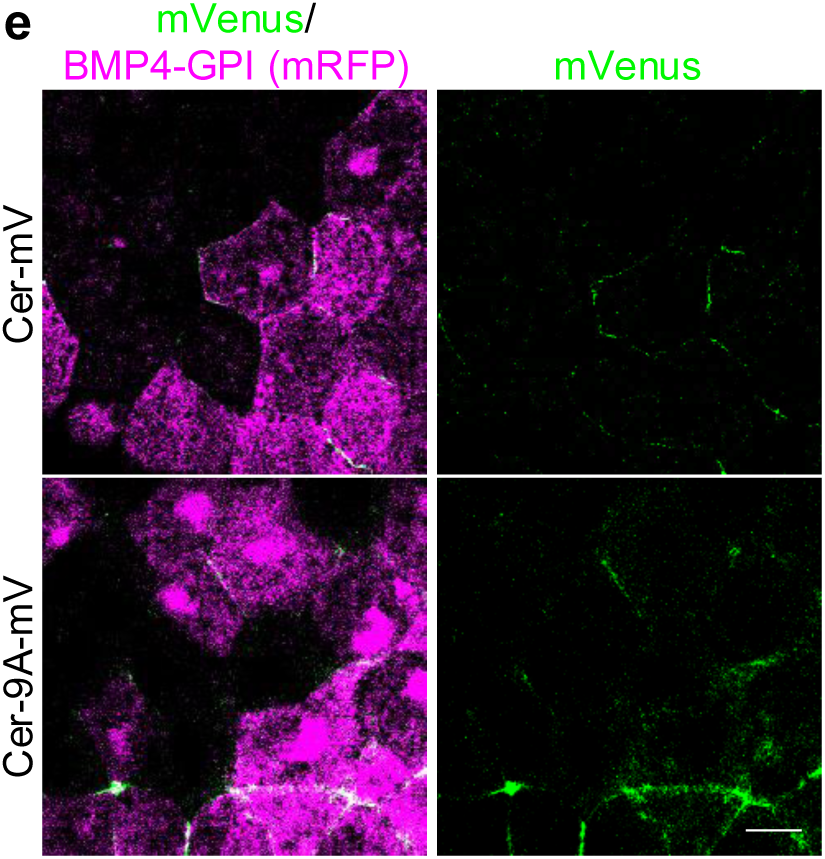
Amino acid sequence and distribution of Cer variants. a, Amino acid sequence of Cer.S (not CerS but Cer.S, indicating *S* gene of Cer (*X. laevis* is an allotetraploid, which has two subgenomes, *L* and *S*)) in *X. laevis*. Basic residues are highlighted in yellow. The residues changed to Ala in Cer variants (Cer-5A or Cer-4A) are colored red. * shows putative cleavage site. Sequence after the arrow is the Cer-short (CerS) (Piccolo et al., 1999). b, Distribution of Cer-mV and CerS-mV. mRNA of Cer-mV or CerS-mV was injected into a blastomere at the 4-cell stage, and specimens were visualized at the gastrula stage. c, Distributions of Cer-mV and CerS-mV on the Ndst1-expressing cells. mRNAs of Cer-mV or CerS-mV and Ndst1 (with the tracer, mRFP) were injected into different blastomeres at the 4-cell stage. d, Cer-mV and Cer-9A-mV were similarly localized to the BMP4-GPI expressing cells. mVenus-tagged Cer variants and BMP4-GPI with mRFP were injected into different blastomeres at the 16-cell stage and the injected embryos were fixed at the early gastrula stage. e, Expansion of distribution range of NAc-HS-binding motif mutants of Frzb and Cer. mRNA of mV-Frzb, mV-Frzb-12A, Cer-mV or Cer-9A-mV was injected into a blastomere at the 4-cell stage, and that of Nanobody (with the tracer, mRFP) was injected into the other two blastomeres as indicated (left). Specimens were observed at the blastula stage. Signal intensity was measured by ImageJ software with the straight-line tool (the width was set to 160 px: around 129 μm). An example of the measurement was described in the left (yellow bold line indicates the measured region). Results were shown in the graphs (dots indicate each value from the measurements; regression curves (cubic spline) were calculated by geom_smooth function in ggplot2 package version 3.3.0 in R version 3.6.3). For Y-axis, signal intensity of mVenus was normalized by each max value of mVenus intensity (normalized intensity). In the upper graphs, the X-axis was shown by actual value of distance from the source (μm). In addition, we have set X axis to the cumulative values of mRFP intensity to the lower graphs because the expression levels of Nanobody (mRFP) were not same among the measured cells. These two graphs similarly indicated that loss of NAc-HS-binding ability expands distribution range of both Frzb and Cer. Amount of mRNA (pg/embryo): Cer-mV/Cer-5A-mV/Cer-4A-mV, 500; CerS-mV, 440 (equivalent mol. of Cer-mV); BMP4-GPI, 520; Ndst1, 500; mRFP, 400. Scale bar represents 20 μm (b-d). 200 μm (e).

**Supplemental Figure 4.**
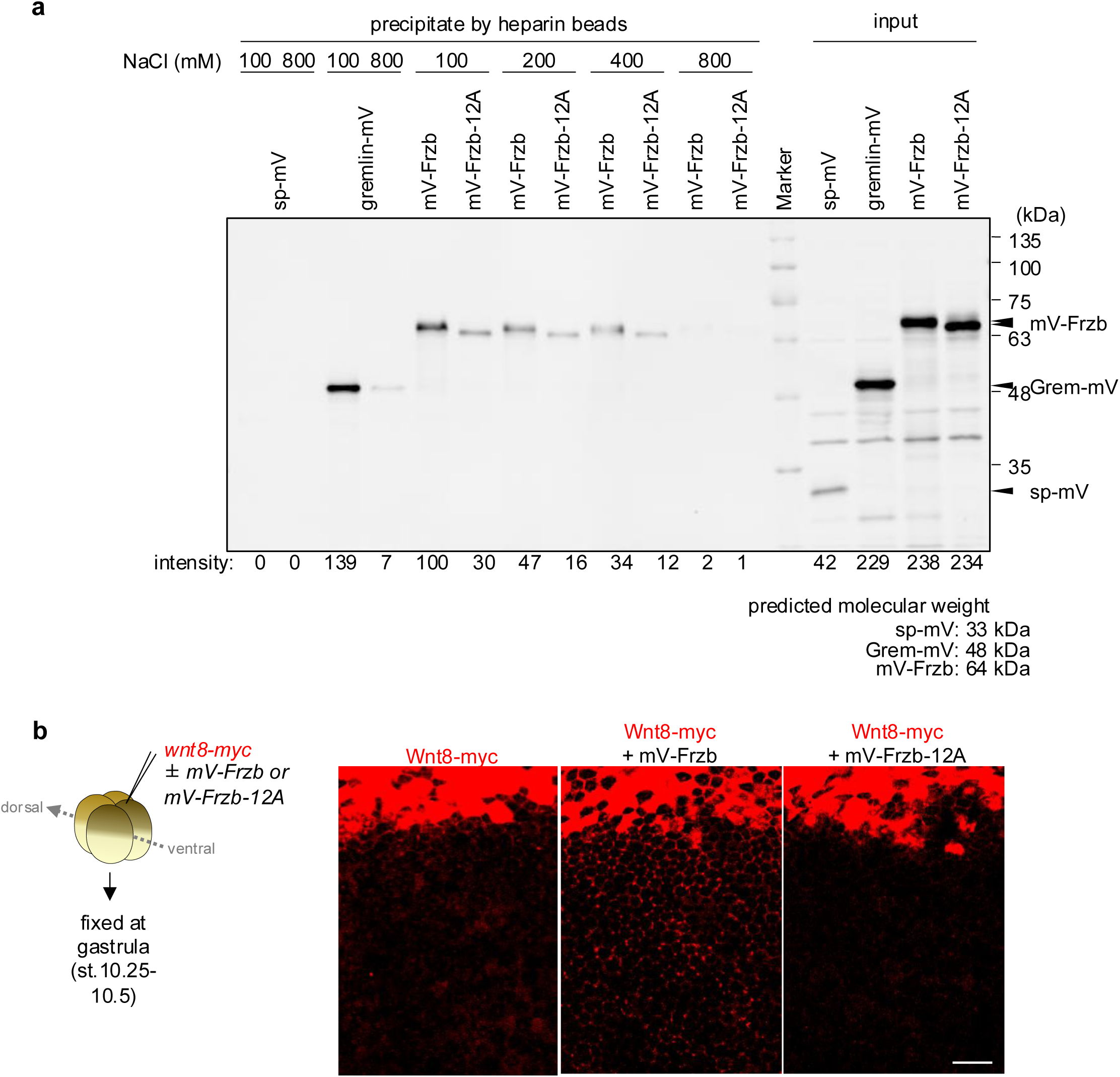
Identification of putative NAc-HS-binding motif of Frzb and Cer. a, Frzb bound to heparin by the putative NAc-HS-binding motif. mRNAs of mV-tagged Frzb variants, grem-mV and sp-mV (signal peptide with mVenus) were injected at the 4-cell stage. Cell extracts from stage 10.5 embryos were incubated with heparin beads, and the beads were washed with lysis buffer containing 100, 200, 400 and 800 mM NaCl. Band quantification was performed with ImageJ. The amount of protein bound to heparin beads, normalized by the mV-Frzb (100 mM NaCl) band, is indicated below each lane. b, Wnt8 distribution with/without Frzb or Frzb-12A. mRNAs of Wnt8-myc and Frzb or Frzb-12A were co-injected into a ventral blastomere at the 4-cell stage. Amount of mRNA (pg/embryo): (a) all mV-tagged mRNA, 200; (b) Wnt8-myc, 374; mV-Frzb or mV-Frzb-12A, 391. Scale bar represents 100 μm (b).

**Supplemental Figure 5.**
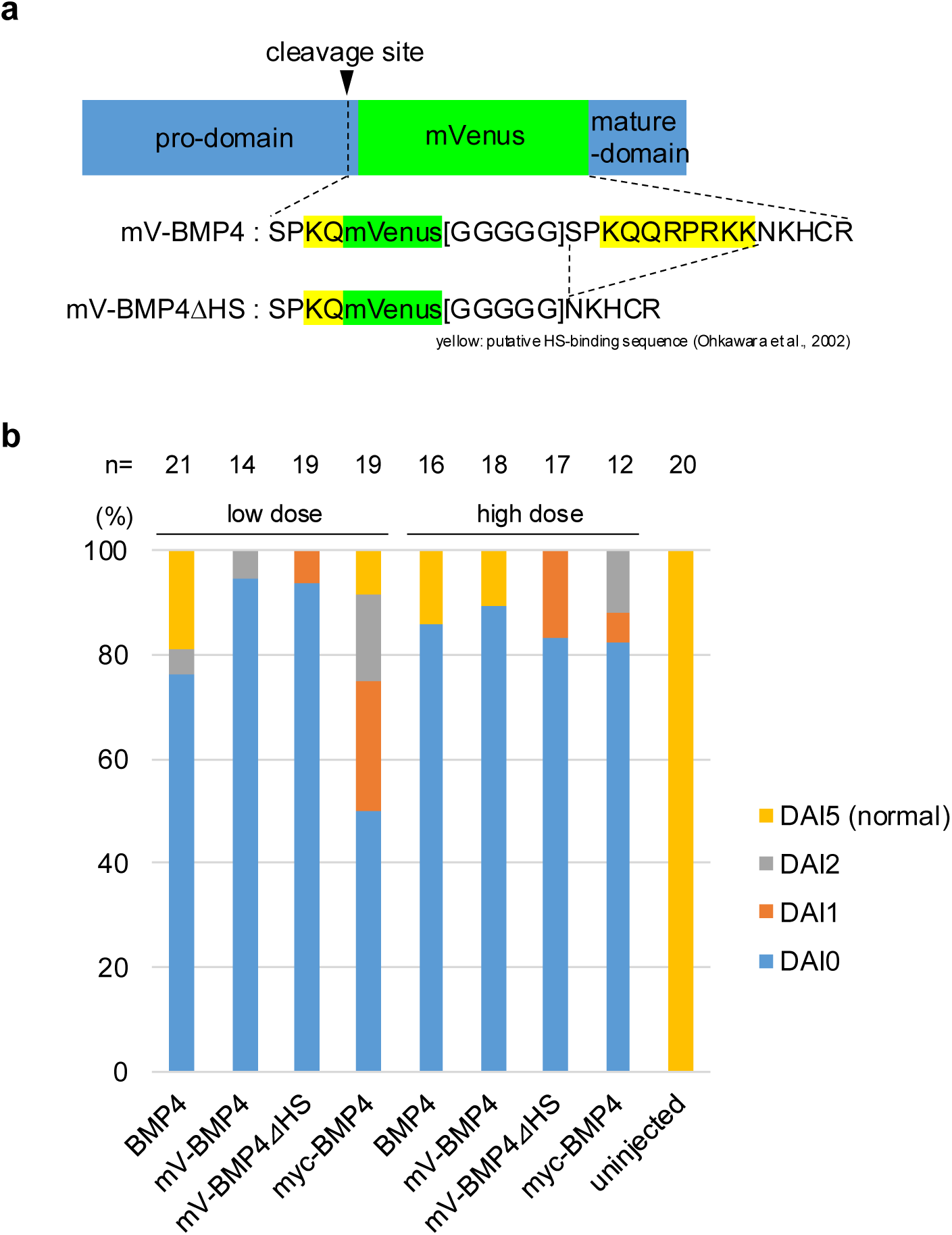
Structure, developmental effect, and distribution of *Xenopus* BMP variants. a, Schematic figure of BMP4 constructs. BMP4 is cleaved into pro-domain and mature domain. mVenus tag (or Myc tag) was inserted into the N-terminal of the mature domain after Gln-290, according to Degnin et al. (2004). The highlighted residues are putative HS binding sequence, proposed by Ohkawara et al. (2003). b, Biological activity of *bmp4* constructs as assayed by ventralization activity. mRNA of each *bmp4* construct was injected into the equatorial region of two dorsal blastomeres near the midline at the 4-cell stage. Phenotypes of embryos were classified with the dorso-anterior index (DAI) (Kao and Elinson, 1988). The grade is numbered from 0 to 10 (0 means no dorso-anterior structures (highly ventralized), and 5 means normal). Percentages of these categories are presented as bar graphs. Total numbers of injected embryos (n) are indicated above.

**Supplemental Figure 6.**
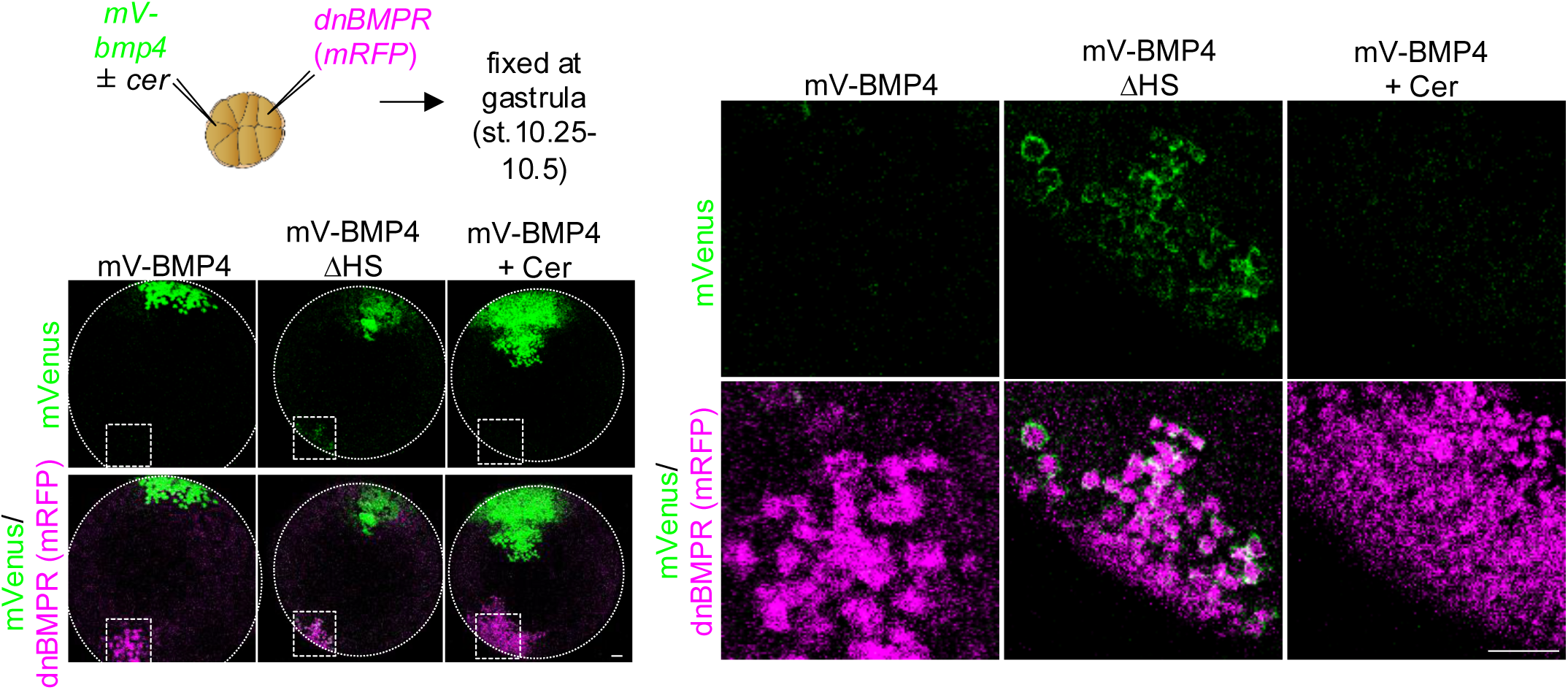
Distribution of BMP4 variants Distribution of BMP4 was restricted by binding to HS. mRNAs of mV-BMP4 variants (see also Supplemental Fig. 6) and dominant-negative BMP receptor (dnBMPR: type I receptor of BMP whose kinase domain was deleted) with a tracer (mRFP) were injected into the animal pole region of different blastomeres (to express these genes mosaically) at the 16-cell stage. Injected embryos were fixed at st. 10.25-10.5. With the deletion of the basic amino acid, the distribution of BMP4 (mV-BMP4ΔHS), even in the far from the source, can be seen on the dnBMPR-expressing cells. Amount of mRNA (pg/embryo): mV-BMP4 variants, 500; dnBMPR, 500; mRFP, 400. Scale bar represents 100 μm.

**Supplemental Figure 7.**
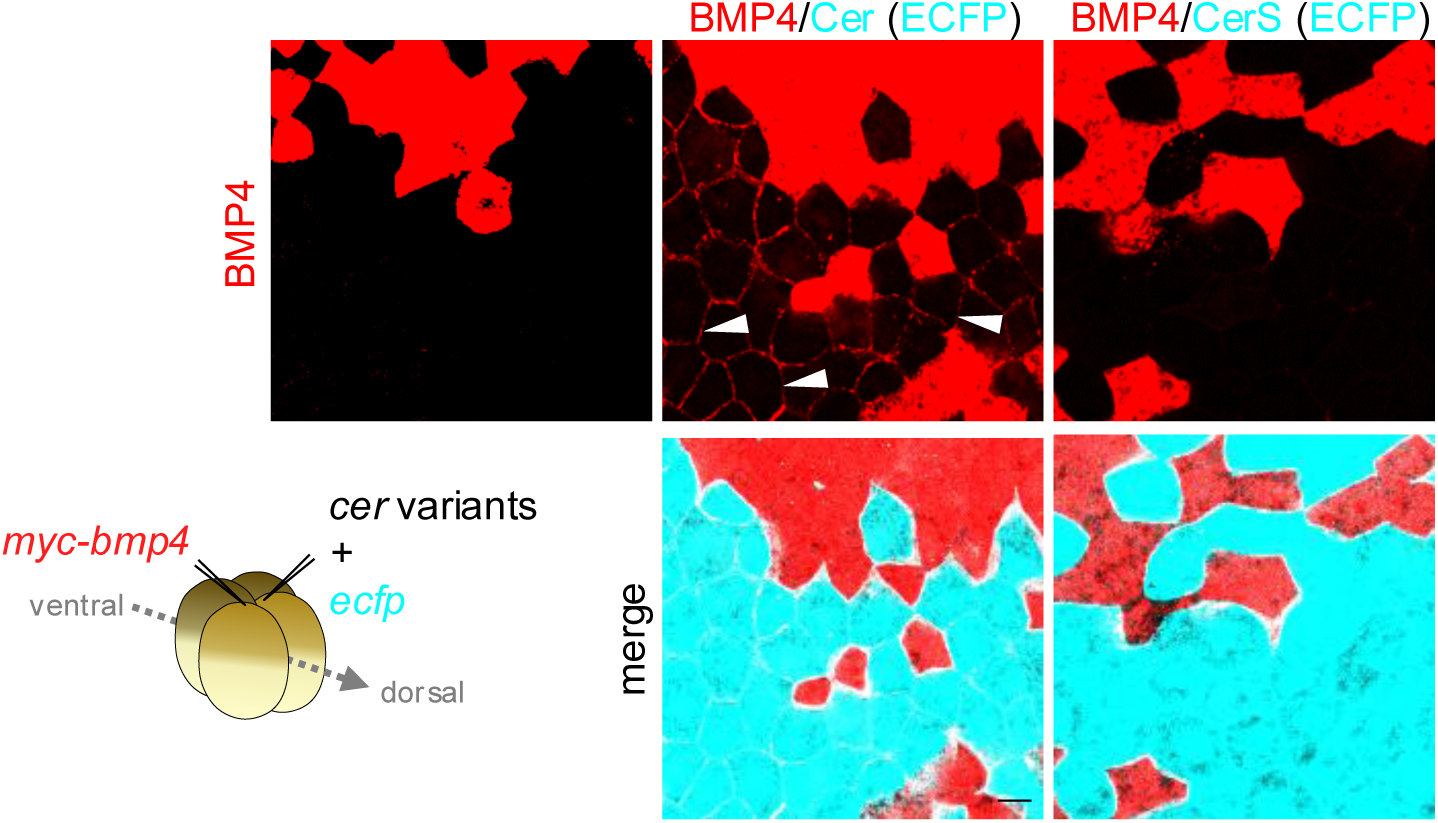
BMP distribution is expanded by Cer even when they are injected into different blastomeres. mRNAs of Myc-BMP4 and Cer variants (with ECFP) were injected into different dorsal blastomeres at the 4-cell stage. Protein distribution of BMP4 was expanded by Cer, but not by CerS. Amount of mRNA (pg/embryo): myc-BMP4, 80; Cer, 61; CerS, 64 (the amount of the antagonists is equivalent mol. to myc-BMP4). Scale bars represent 30 μm.

**Supplemental Figure 8.**
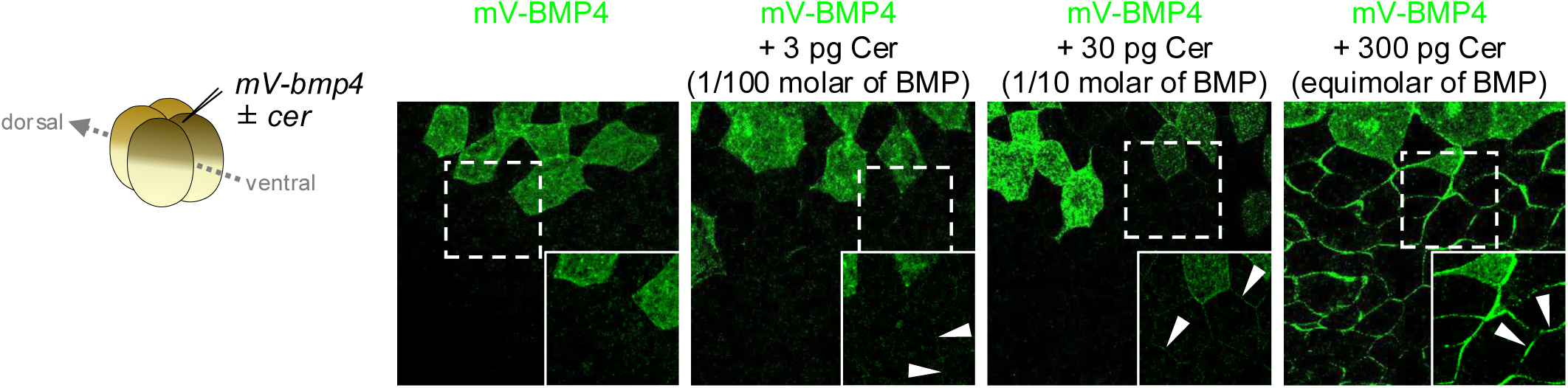
Cer accumulates BMP4 in the intercellular region. Cer accumulated BMP4 on the cell surface, regardless of Cer concentration. Various amount of Cer mRNA was injected (as described in the figure) with 500 pg mV-BMP4 mRNA. Scale bar represents 30 μm.

**Supplemental Figure 9.**
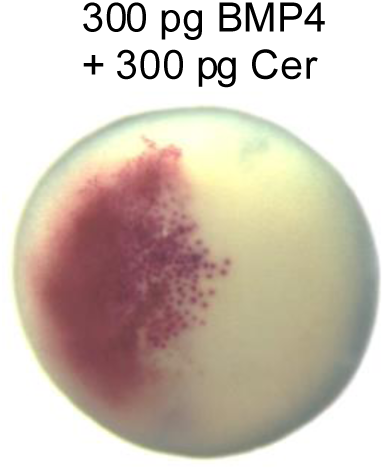
B**M**P **activity in the antagonist-coinjected embryos** High amount (300 pg) of Cer mRNA injection inhibits BMP signaling. mRNAs of BMP4 and nβ-gal (tracer) with Cer were injected into a dorsal blastomere at the 4-cell stage, and the specimens were fixed at the gastrula stage (st.10.5). Source cells were colored red by Red-gal.

---

**Supplemental Table 1.**
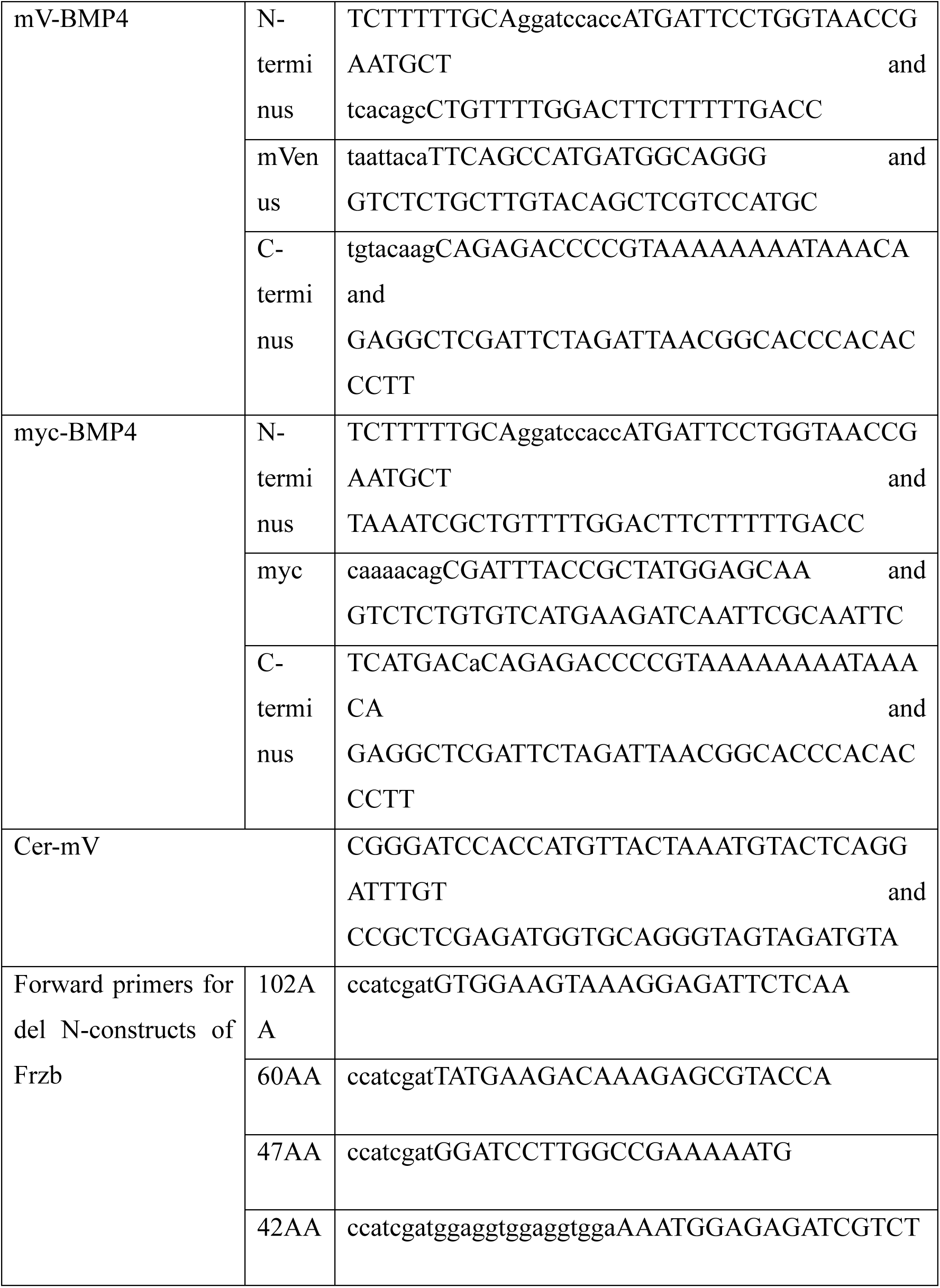
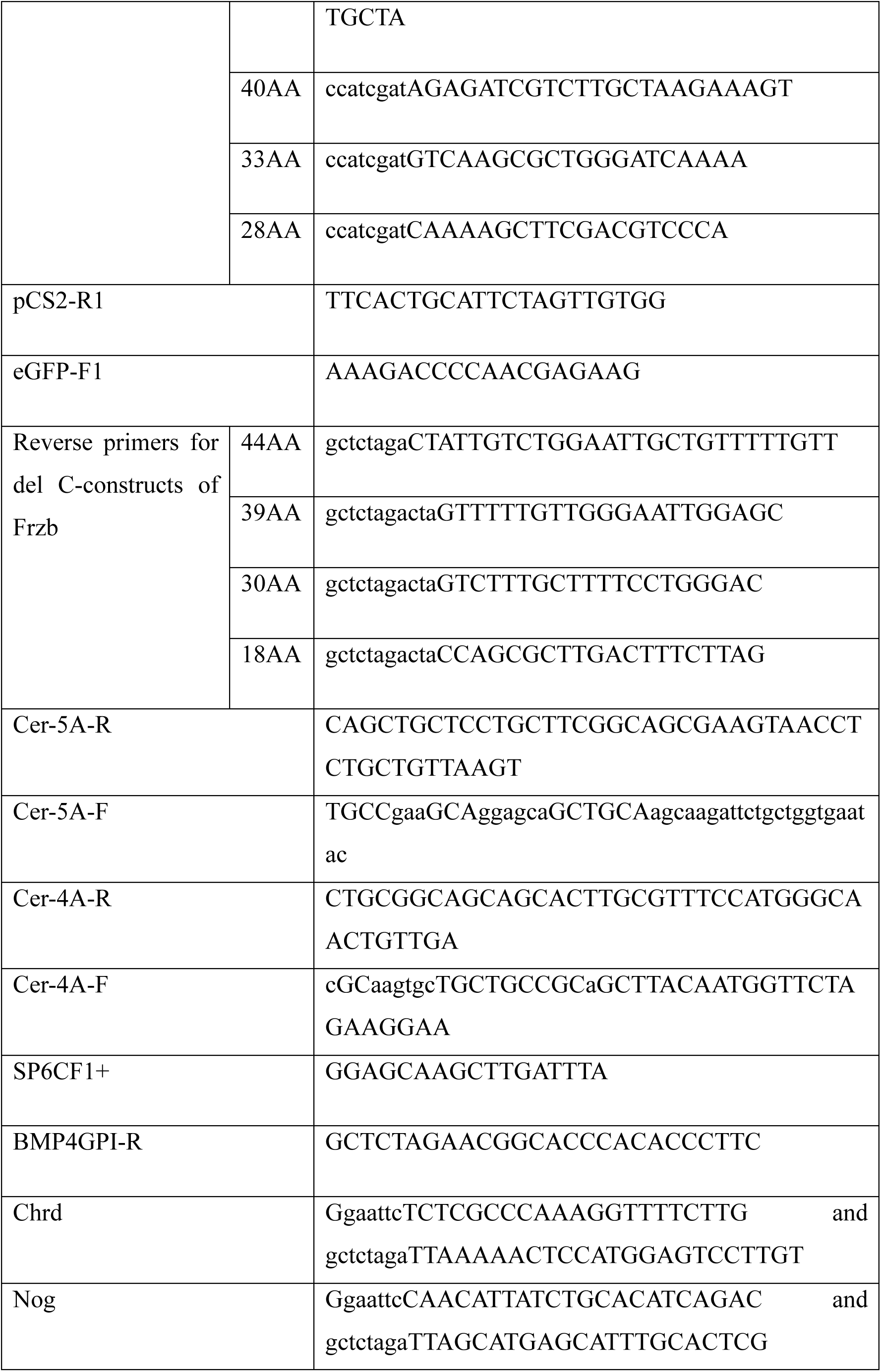
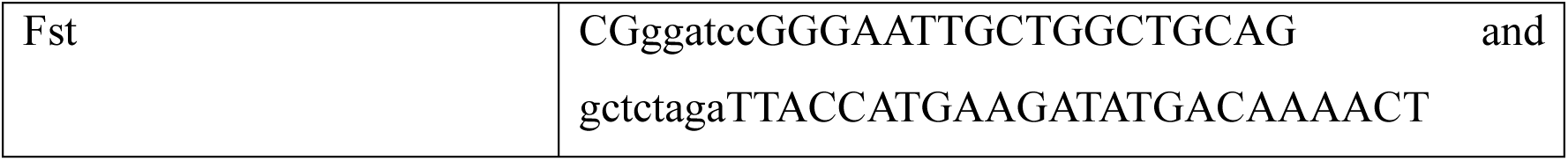
Primers used in this study.

## Notes

### Competing Interest Statement

The authors have declared no competing interest.

